# The TAZ2 domain of CBP/p300 directs acetylation towards H3K27 within chromatin

**DOI:** 10.1101/2020.07.21.214338

**Authors:** Thomas W. Sheahan, Viktoria Major, Kimberly M. Webb, Elana Bryan, Philipp Voigt

## Abstract

The closely related acetyltransferases CBP and p300 are key regulators of gene expression in metazoans. CBP/p300 acetylate several specific lysine residues within nucleosomes, including histone H3 lysine 27 (H3K27), a hallmark of active enhancers and promoters. However, it has remained largely unclear how specificity of CBP/p300 towards H3K27 is achieved. Here we show that the TAZ2 domain of CBP is required for efficient acetylation of H3K27, while curbing activity towards other lysine residues within nucleosomes. We find that TAZ2 is a sequence-independent DNA binding module, promoting interaction between CBP and nucleosomes, thereby enhancing enzymatic activity and regulating substrate specificity of CBP. TAZ2 is further required to stabilize CBP binding to chromatin in mouse embryonic stem cells, facilitating specificity towards H3K27 and modulating gene expression. These findings reveal a crucial role of TAZ2 in regulating H3K27ac, while highlighting the importance of correct site-specific acetylation for proper regulation of gene expression.

## Introduction

In eukaryotes, transcription is regulated through a range of posttranslational modifications to the histone proteins that package DNA into chromatin. Histone acetylation is thought to play key roles in this process by directly regulating accessibility to the underlying DNA and by recruiting effector proteins featuring acetyl-lysine binding bromodomains (Filippakopoulos and Knapp, 2012; Zentner and Henikoff, 2013). All four core histone proteins can be acetylated at numerous lysine residues by a range of histone acetyltransferase (HAT) enzymes (Barnes et al., 2019; Marmorstein and Zhou, 2014). Histone acetylation marks at different residues commonly co-occur in combinatorial fashion, colocalizing at regulatory regions of actively transcribed genes (Birney et al., 2007; Heintzman et al., 2007; Wang et al., 2008). Indeed, acetylation of histone H3 lysine 9 (H3K9), H3K18, H3K27, and of several residues within the N-terminal tail of histone H4 have all been shown to correlate with active promoters and enhancers (Creyghton et al., 2010; Karmodiya et al., 2012; Rada-Iglesias et al., 2011; Wang et al., 2009; Wang et al., 2008; Zhang et al., 2020) Despite this extensive cooccurrence, specific functions have been demonstrated for individual acetylation marks, including H4K16ac in regulating interactions between neighbouring nucleosomes (Shogren-Knaak et al., 2006) and H3K9ac in facilitating recruitment of the basal transcription factor TFIID (Vermeulen et al., 2007). These observations suggest cooperative rather than redundant functions for specific acetylation marks, highlighting the importance of HAT substrate specificity for proper regulation of transcription. While it has been well established that most HATs target only a small subset or even unique histone lysine residues, the mechanisms by which individual HATs achieve specificity for their cognate substrates remain largely elusive.

Genome-wide mapping studies have attributed a particular importance to acetylation of H3K27 (H3K27ac), and this modification is widely used as a marker to distinguish active from inactive regulatory elements, particularly in the case of enhancers (Creyghton et al., 2010; Rada-Iglesias et al., 2011). In addition to supporting recruitment of specific bromodomain proteins including components of the super elongation complex (SEC) and the BAF complex (Jefimov et al., 2018), H3K27ac prevents placement of H3K27 methylation by Polycomb Repressive Complex 2 (PRC2), thereby antagonizing gene silencing (Pasini et al., 2010; Tie et al., 2009). H3K27ac is generated by the enzymes CBP and p300, with H3K27ac essentially absent upon inhibition or combined knockout of CBP and p300 in mammalian cells (Jin et al., 2011; Weinert et al., 2018). CBP and p300 (together referred to as CBP/p300) are homologous HAT enzymes that are highly conserved among metazoans and some closely related unicellular organisms (Marmorstein and Zhou, 2014; Sebé-Pedrós et al., 2011). Both CBP and p300 are essential for development in mice (Oike et al., 1999; Tanaka et al., 1997; Yao et al., 1998) and mutations in either gene are associated with the rare developmental disorders Rubinstein-Taybi syndrome and Menke-Hennekam syndrome in humans (Hennekam, 2006; Menke et al., 2016; Petrij et al., 1995). CBP/p300 display relatively narrow substrate specificity towards histones and in addition to H3K27ac also contribute to acetylation of H3K18 and residues within the N-terminal tail of histone H2B such as H2BK5 and H2BK12 (Jin et al., 2011; Weinert et al., 2018). However, it remains unclear how these physiological substrate preferences are specified mechanistically.

CBP/p300 is a large multi-domain protein organized around the catalytic HAT domain (Figure 1A). The HAT domain along with the bromo, PHD, and RING domains form the catalytically active CBP core (Delvecchio et al., 2013; Park et al., 2017). However, despite being sufficient for catalytic activity, the CBP core fails to recapitulate the site specificity of full-length CBP/p300 observed *in vivo* (Bannister and Kouzarides, 1996; Zhang et al., 2018), suggesting that domains or motifs outside of the catalytic core determine substrate specificity of CBP/p300. Moreover, CBP/p300 are less active and more site-selective towards histones assembled into nucleosomes than free histones (An and Roeder, 2003; Kraus et al., 1999), suggesting that CBP/p300 activity and specificity also depends on the chromatin context of their histone substrates. Domains within CBP/p300 that interact with features of nucleosomes could therefore play a role in regulating enzymatic activity and determining substrate specificity. Recently, the ZZ domain was shown to interact with the N terminus of histone H3, thereby specifically promoting acetylation of H3K27 in nucleosome core particle (NCP) substrates (Zhang et al., 2018). However, it remains unclear whether other domains further contribute to regulation of catalytic activity and specificity of CBP/p300.

**Figure 1.**
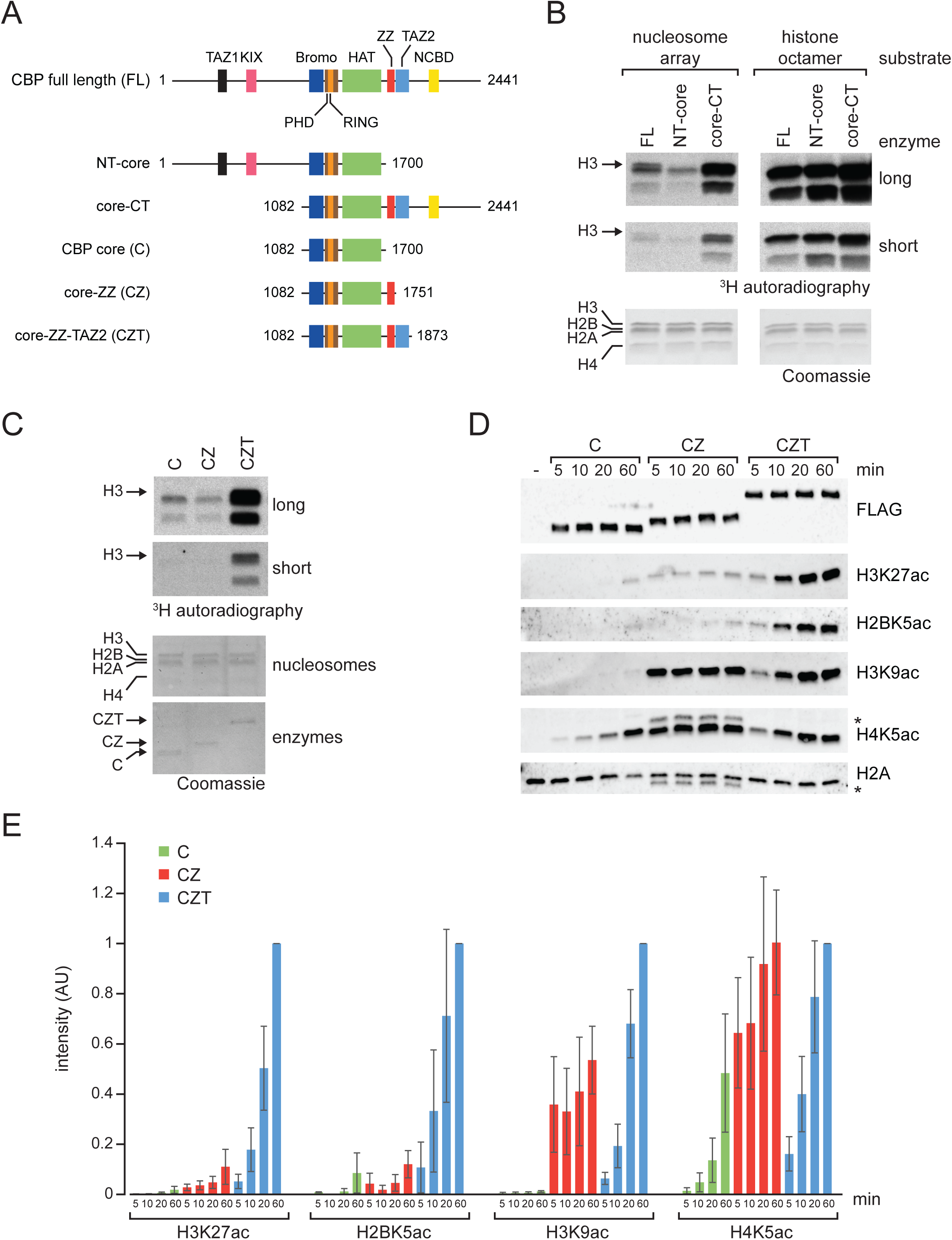
The CBP TAZ2 domain is required for efficient acetylation of H3K27 *in vitro*. (**A**) Schematic representation of the domain architecture of full-length (FL) mouse CBP and CBP truncation proteins, showing unstructured regions (line) and structured domains (boxes), with numbers representing amino acid positions in full-length mouse CBP. TAZ1, transcription adaptor zinc finger domain 1; KIX, kinase-inducible domain (KID)-interacting domain; Bromo, bromodomain; PHD, plant homeodomain zinc finger; RING, really interesting new gene zinc finger; HAT, histone acetyltransferase domain; ZZ, ZZ-type zinc finger; TAZ2, transcription adaptor zinc finger domain 2; NCBD, nuclear co-activator binding domain. (**B**) HAT assays using CBP FL, NT-core, and core-CT constructs with nucleosome arrays (left panels) or histone octamer (right panels) as substrates. Reactions were performed for 20 min. Coomassie staining of membrane shows equal loading of histone proteins, with individual histones indicated, and ^3^H autoradiographs show long and short autoradiograph exposures. Arrows indicate histone H3. (**C**) HAT assays using CBP core (C), core-ZZ (CZ), and core-ZZ-TAZ2 (CZT) constructs with nucleosome array substrates. Coomassie staining shows loading of histones and enzymes, and ^3^H autoradiographs show long and short exposures. Assays shown in (B,C) are representative of three independent experiments each. (**D**) Western blot analysis of time course HAT assays with no enzyme (-), C, CZ, or CZT enzymes with nucleosome array substrates. Reactions were analyzed using antibodies against FLAG, H3K27ac, H2BK5ac, H3K9ac, H4K5ac, and total H2A. Asterisks indicate non-specific bands present in CZ reactions (see Figure S1D). (**E**) Quantification of HAT assays shown in (D). Signal for each antibody is normalized to signal obtained with CZT at 60 min. Data shown represents the mean and SEM of three independent experiments. See also Figure S1.

Here, we show that the TAZ2 domain of CBP is required for efficient and targeted acetylation of H3K27. The TAZ2 domain possesses a novel DNA binding activity that stabilizes CBP binding to chromatin and regulates CBP substrate specificity *in vitro* and *in vivo*. These findings suggest that the TAZ2 domain is an integral part of the CBP histone acetylation machinery, allowing precise regulation of gene expression through site-specific acetylation of histones.

## Results

### The TAZ2 domain of CBP is required for efficient acetylation of H3K27 *in vitro*

To determine which domains outside the catalytic core of CBP contribute to histone acetylation activity and specificity, a domain-mapping approach was utilized. We expressed and purified a panel of truncated CBP proteins (Figure 1A, S1B) and tested their activity in HAT assays, using either histone octamers or nucleosome arrays (Figure S1A) as substrates. Full-length CBP was able to acetylate all four core histones in nucleosome arrays, with the highest activity observed for histones H3 and H2B (Figure 1B, left panels). Whereas this acetylation pattern was unaffected by deletion of the N terminus, which contains the transcriptional activator zinc finger (TAZ) domain 1 and the kinase-inducible CREB interaction region (KIX), deletion of the C terminus abrogated acetylation of histone H3 in nucleosome arrays (Figure 1B, left panels). Importantly, all three constructs were able to acetylate histone H3 when using histone octamers as substrates (Figure 1B, right panels), suggesting that regions in the C terminus of CBP are required for acetylation of histone H3 specifically in the context of physiologically relevant chromatin substrates such as nucleosome arrays.

The C terminus of CBP features a ZZ-type zinc finger and a second TAZ-type zinc finger called TAZ2 immediately downstream of the CBP core, followed by a nuclear coactivator binding domain (NCBD). To determine which of these domains are necessary for acetylation of histone H3 in nucleosome arrays, further truncated proteins were generated comprising only the catalytic core of CBP, or the core together with the neighbouring ZZ domain (CZ) or together with both the ZZ and TAZ2 domains (CZT) (Figure 1A, S1C). The isolated catalytic core was unable to efficiently acetylate histone H3, whilst inclusion of the ZZ domain led to a modest but reproducible increase in activity towards histone H3 (Figure 1C), consistent with previous studies reporting that the ZZ domain of CBP/p300 is important for H3 acetylation (Bannister and Kouzarides, 1996; Zhang et al., 2018). Addition of the TAZ2 domain, however, was both necessary and sufficient to achieve robust histone H3 acetylation, while also increasing activity towards the other core histones (Figure 1C). These findings indicate that the TAZ2 domain of CBP is required for efficient acetylation of histone H3 in chromatin substrates.

To understand whether the ZZ and TAZ2 domains influence acetylation of specific lysine residues, we carried out time course HAT assays with nucleosome arrays and analyzed acetylation of H3K27 and H2BK5, two physiological targets of CBP, and of H3K9 and H4K5, acetylation of which is independent of CBP *in vivo*. Consistent with the radiolabel-based HAT assays reflecting overall catalytic activity (Figure 1C), the isolated CBP core only weakly acetylated H3K27 and H2BK5, primarily generating off-target H4K5ac (Figure 1D, E). The CZ construct displayed markedly increased activity towards H3K9 and H4K5, but only modestly enhanced activity towards the CBP targets H3K27 and H2BK5 (Figure 1D, E). In contrast, the CZT construct efficiently acetylated the CBP target sites H3K27 and H2BK5 (Figure 1D), generating ten-fold higher levels of H3K27ac and H2BK5ac while exhibiting lower activity towards H3K9 and H4K5 than CZ (Figure 1D, E). These results suggest that the TAZ2 domain specifies CBP activity towards key physiological substrates such as H3K27 and H2BK5, while curbing off-target activity.

### The TAZ2 domain of CBP is a sequence-independent DNA binding module

Having established that the TAZ2 domain is required for efficient acetylation of H3K27, we next sought to determine how it functions to regulate the substrate specificity of CBP. The TAZ2 domain of CBP/p300 is a zinc finger comprising three HCCC-type zinc-coordinating clusters and four alpha-helices (Figure S2A, B) and has been shown to mediate protein-protein interactions with partners such as p53, STAT1 and C/EBP (Bhaumik et al., 2014; Jenkins et al., 2009; Wojciak et al., 2009). However, these interaction partners are absent from the reconstituted system used here, suggesting direct interaction of the TAZ2 domain with either histones or DNA present in the HAT assays. Sequence alignment of TAZ2 domains from CBP/p300 orthologues shows high conservation even in distantly related species (Figure S2A), which not only covers key structural residues but also extends to multiple positively charged lysine and arginine residues. Moreover, calculation of the surface charge based on a previously solved crystal structure (Miller et al., 2009) indicates that the surface of the TAZ2 domain is highly positively charged (Figure S2B, C). We therefore hypothesized that the TAZ2 domain could interact with negatively charged molecules such as DNA, thereby regulating CBP substrate specificity.

To test whether TAZ2 interacts with DNA, we carried out DNA binding assays in which purified recombinant TAZ2 (Figure S2D) was incubated with streptavidin beads alone or coated with a biotinylated 147-bp DNA corresponding to the 601 nucleosome positioning sequence (Lowary and Widom, 1998). Whilst the ZZ domain failed to bind in either condition, the TAZ2 domain was specifically bound only in the presence of DNA (Figure 2A). Specific interaction of TAZ2 with DNA was also observed in electrophoretic mobility shift assays (EMSAs), showing a concentration-dependent band shift of 601 DNA, whereas the ZZ domain was unable to bind DNA (Figure 2B). To test whether TAZ2 exhibits sequence preferences or binds DNA independently of its sequence, further EMSAs were carried out using four 29-bp DNA probes with randomly generated sequences of differing GC content. The TAZ2 domain bound to these probes with similar affinity (Figure 2C), suggesting that TAZ2 binds DNA in a sequence-independent manner.

**Figure 2.**
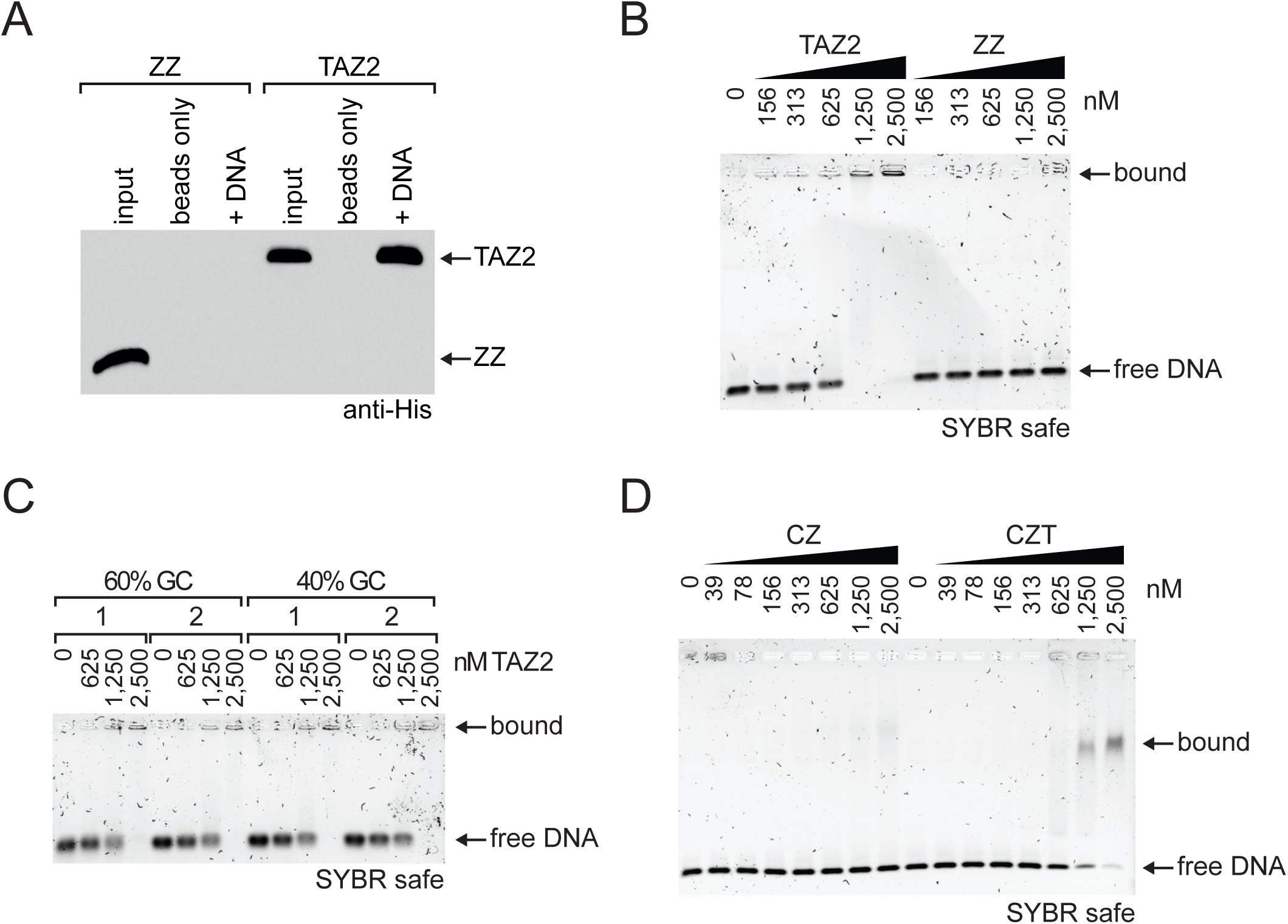
The CBP TAZ2 domain is a sequence-independent DNA binding module. (**A**) DNA pulldown assay with isolated ZZ or TAZ2 domain and either streptavidin beads alone (beads only) or streptavidin beads coated with biotinylated 147-bp 601 DNA (+ DNA). Pulldown assays were analyzed by western blot using anti-His antibody. Pulldown shown is representative of three independent pulldown experiments. (**B–D**) EMSA assays for DNA binding of CBP constructs. DNA is visualized by agarose gel electrophoresis and SYBRsafe staining. Data shown is representative of three independent EMSA experiments. (B) Titration of isolated TAZ2 or ZZ domain with 147-bp 601 DNA; (C) titrations of TAZ2 domain with 29-bp dsDNAs of differing GC content and sequence; (D) titrations of CBP core-ZZ (CZ) or core-ZZ-TAZ2 (CZT) constructs with 147-bp DNA. See also Figure S2.

To confirm that the DNA binding activity of TAZ2 is retained in the context of a catalytically competent CBP protein, EMSA experiments were carried out using CZ and CZT proteins. Whilst CZ was unable to bind the 147-bp DNA probe, inclusion of the TAZ2 domain was sufficient to achieve DNA binding (Figure 2D). Together, these results show that the TAZ2 domain of CBP binds DNA in a sequence-independent manner and can mediate interactions between an enzymatically competent CBP protein and free DNA.

### The TAZ2 domain mediates interactions between CBP and nucleosomes

Given that the TAZ2 domain of CBP binds free DNA, we next asked whether this domain also mediates interaction with DNA in the context of nucleosomes, the physiological substrate of CBP. To this end, EMSA experiments were performed with isolated TAZ2 or ZZ domains and nucleosome core particles (NCPs) assembled with either a minimal 147-bp 601 nucleosome positioning sequence or a longer 209-bp sequence with additional symmetric 31-bp DNA overhangs. Whereas the ZZ domain did not exhibit binding to NCPs at any of the concentrations tested, the TAZ2 domain robustly interacted with NCPs irrespective of the presence of linker DNA (Figure 3A, B), suggesting that TAZ2 can bind to nucleosomal DNA as well as to free DNA.

**Figure 3.**
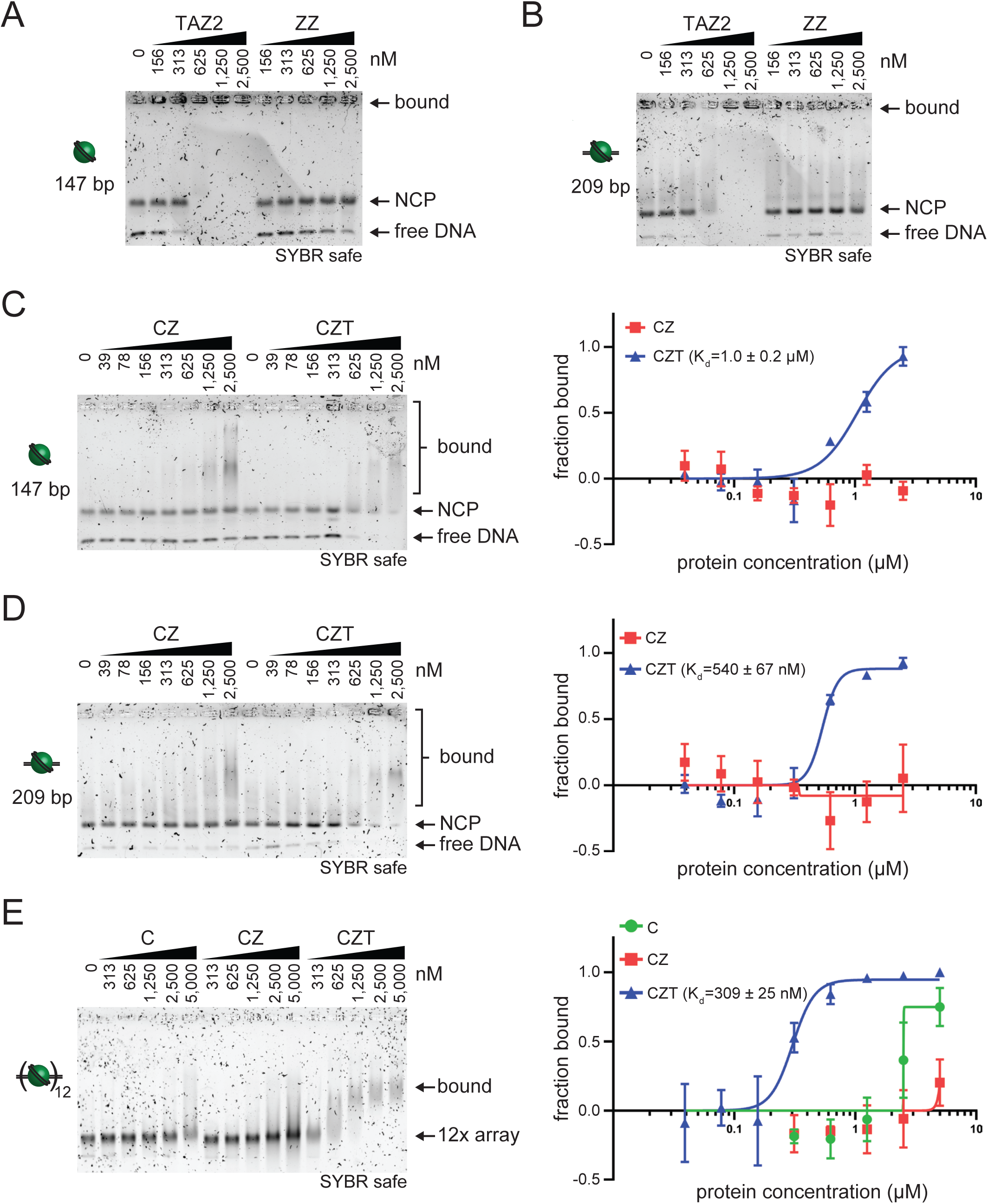
The TAZ2 domain mediates interactions between CBP and nucleosomes. EMSA assays with nucleosomal DNA templates, analyzed by agarose gel electrophoresis and SYBRsafe staining. Data shown are representative of three independent experiments each. (**A**), (**B**) Binding reactions for isolated TAZ2 or ZZ domain with 147-bp (A) or 209-bp (B) nucleosome core particles (NCP). (**C**), (**D**) Binding reactions for CZ and CZT constructs with 147-bp (C) or 209-bp (D) NCPs. (**E**) Binding reactions for CBP core (C), CZ or CZT constructs with 12x nucleosome arrays. (C-E), Right: quantitative analysis of EMSA data, yielding apparent K_d_ values. Plots show data mean and SEM from three independent experiments, measured by depletion of free nucleosome array band.

To determine whether nucleosome binding was also observed in the context of enzymatically competent CBP proteins, further EMSA experiments were carried out using CZ and CZT proteins and NCPs. Whereas CZ bound NCPs with only low affinity, inclusion of the TAZ2 domain strongly increased affinity towards NCPs (Figure 3C, D), binding 147-bp and 209-bp NCPs with a K_d_ of 1.0 μM and 0.54 µM, respectively. By contrast, the reported K_d_ for interactions between the ZZ domain and histone H3 is 8.8 µM (Zhang et al., 2018), suggesting that the TAZ2 domain mediates interactions with nucleosomes with considerably higher affinity than the ZZ domain. To test whether TAZ2 is also important for interactions with more complex chromatin templates, EMSAs were carried out with nucleosome arrays and CBP core, CZ, and CZT proteins. Whilst only weak binding was observed for CBP core and CZ, CZT robustly bound nucleosome arrays with a K_d_ of 309 nM (Figure 3E). Together, these results indicate that the TAZ2 domain is both necessary and sufficient to mediate high-affinity interactions between CBP and chromatin substrates.

### TAZ2–nucleosome interactions regulate CBP enzymatic activity and substrate specificity

Having established that the TAZ2 domain is crucial for interactions between CBP and nucleosome substrates, we next sought to determine the consequences of these interactions for CBP enzymatic activity. As shown in Figure 1C, the TAZ2 domain enhanced overall enzymatic activity of the CBP core. We reasoned that if the increased activity was mediated by TAZ2-dependent interactions with DNA in nucleosomal arrays, addition of free DNA would compete with TAZ2-nucleosome interactions and thus reduce activity of CBP. Indeed, addition of nucleosome-free DNA curtailed activity of full-length CBP towards nucleosomal arrays, whereas activity of a core construct lacking the TAZ2 domain was unaffected (Figure 4A). Sensitivity towards free DNA was recovered when the core was fused to the C terminus of CBP containing the TAZ2 domain (Figure 4A). These findings indicate that the DNA binding properties of TAZ2 mediate stimulation of CBP activity.

**Figure 4.**
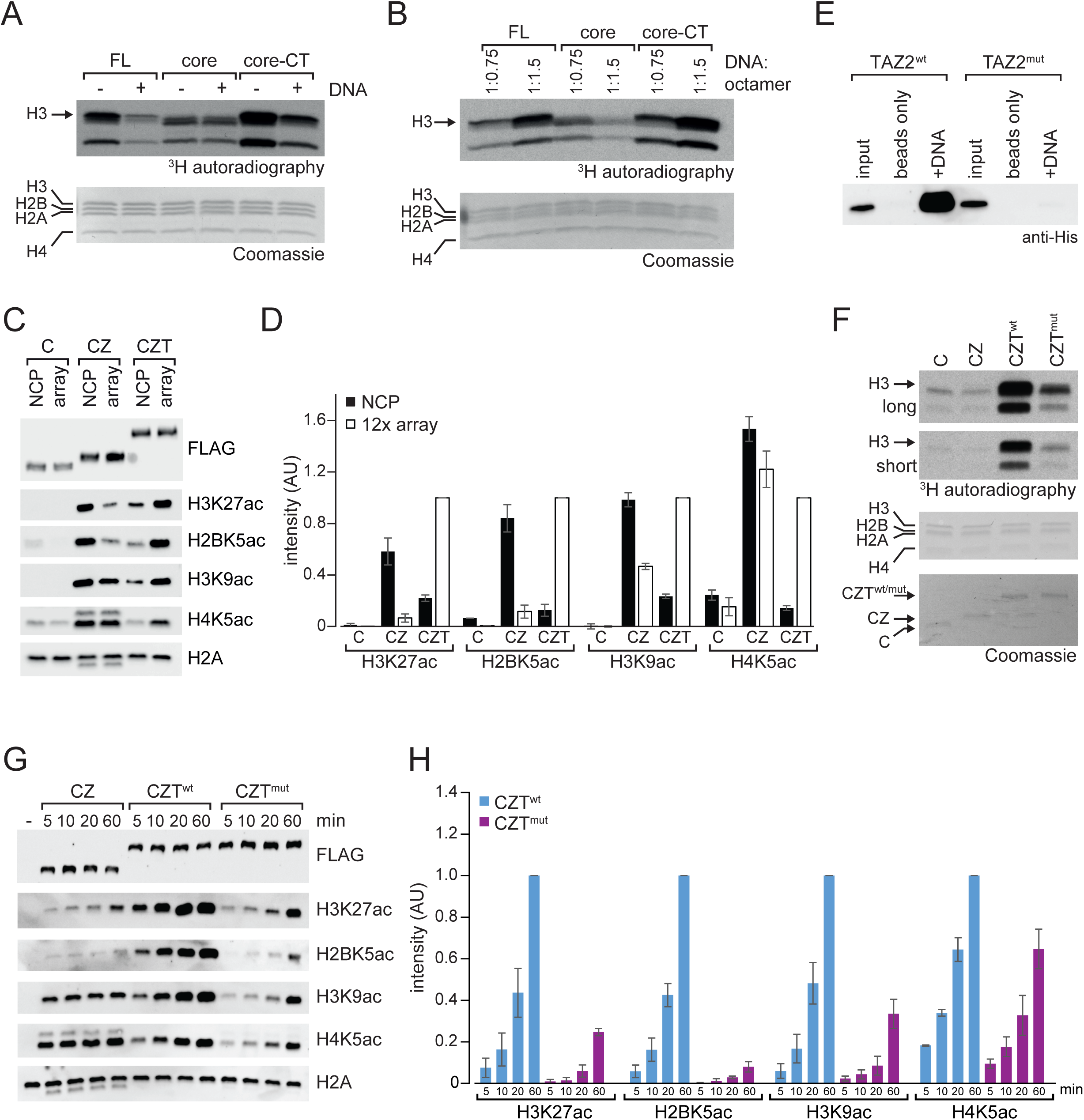
TAZ2–nucleosome interactions regulate CBP enzymatic activity and substrate specificity. **(A)** HAT assays using CBP FL, p300 core, and core-CT with nucleosome arrays in the absence (-) or presence (+) of competitor free DNA. Coomassie staining of membrane shows equal loading of histone proteins, with individual histones indicated, and ^3^H shows autoradiograph exposure. Arrows indicate histone H3. **(B)** HAT assays using CBP FL, p300 core, and core-CT constructs with nucleosome arrays reconstituted with different ratios of DNA:histone octamer. Reactions were performed and evaluated as in (A). (**C**) Western blot analysis of HAT assays with CBP core (C), CZ, and CZT constructs and 147-bp NCP or 12x nucleosome arrays as substrates. (**D**) Quantification of (C). Data shown represents mean and SEM of three experiments normalized to CZT reactions with 12x nucleosome array. Black and white bars represent reactions with NCPs and 12x nucleosome arrays, respectively. (**E**) DNA pulldown assay with TAZ2^wt^ and TAZ2^mut^ domains and either streptavidin beads alone (beads only) or coated with biotinylated 147-bp 601 DNA (+DNA). Pulldown assays were analysed by western blot using anti-His antibody. Assay shown is representative of three independent experiments. (**F**) HAT assays using CBP core, CZ, CZT^wt^, or CZT^mut^ constructs with nucleosome arrays as substrate. Coomassie staining verifies equal loading of histone proteins, with individual histones indicated, and ^3^H autradiographs show long and short exposures. Arrow highlights histone H3. (**G**) Time course HAT assays with no enzyme (-), CZ, CZT^wt^, or CZT^mut^ enzymes and nucleosome arrays as substrates. Reactions were analyzed by Western blot using antibodies against FLAG, H3K27ac, H2BK5ac, H3K9ac, H4K5ac, or total H2A. (**H**) Quantification of HAT assays shown in (G). Signal for each antibody is normalized to CZT^wt^ reaction at 60 min. Data shown represents mean and SEM of three independent experiments. See also Figure S3.

We further reasoned that increasing the density of substrate lysines on DNA would result in more efficient histone acetylation by TAZ2-containing CBP constructs. To test this hypothesis, we prepared nucleosome arrays with increasing levels of histone octamer saturation (Figure S1A). As expected, full-length CBP exhibited higher activity towards substrates with higher numbers of nucleosomes–and thus substrate sites– per DNA molecule, whereas the isolated catalytic core exhibited comparable activity towards both substrates (Figure 4B). Reintroducing the TAZ2 domain by fusing the core to the C terminus of CBP recovered the increased activity on more densely packed nucleosomal arrays (Figure 4B). The TAZ2 domain was necessary and sufficient for enhanced acetylation, whereas addition of the ZZ domain alone did not increase activity with increased nucleosome density (Figure S3A). These results further support that the TAZ2 domain enhances association with nucleosome substrates via DNA binding, leading to higher CBP activity with higher histone:DNA ratios.

To test the role of DNA binding in TAZ2-mediated substrate specificity of CBP, we asked whether TAZ2-containing CBP constructs exhibited differential activity towards specific lysines in NCPs and nucleosome arrays. Whilst CBP core displayed negligible activity towards either substrate, inclusion of the ZZ domain resulted in robust activity towards H3K27, H2BK5, H3K9 and H4K5 in NCP substrates (Figure 4C, D). This observation shows that the ZZ domain enhances acetylation towards multiple lysines in NCPs, not only towards H3K27 as reported previously (Zhang et al., 2018). Interestingly, assembly of nucleosomes into chromatin arrays diminished CZ activity in a site-specific manner, resulting in an approximately ten-fold loss of activity towards H3K27 and H2BK5 but only marginal decrease in activity towards H3K9 and H4K5 when compared to NCPs (Figure 4C, D). These observations indicate that ZZ-mediated binding of the N-terminal tail of histone H3 is sufficient to enable transient interaction of CBP with nucleosome substrates, allowing acetylation of all sites tested in NCPs. However, for the more physiological nucleosomal array substrates, inclusion of the ZZ domains only supports efficient acetylation of H3K9 and H4K5 but not of the major *in vivo* CBP targets H3K27 and H2BK5.

By contrast, a construct containing both ZZ and TAZ2 domains (CZT) acetylated nucleosomal arrays more efficiently than NCPs (Figure 4C, D). This behaviour is consistent with the DNA binding activity of TAZ2 markedly enhancing affinity of CZT towards nucleosomal substrates compared to CZ (see Figure 3). Both CZ and CZT constructs would initially engage with NCPs and nucleosome arrays by random collision. For CZ, collision would result in only transient interaction, allowing rapid dissociation and re-engagement with the next substrate, thus favouring activity towards the more abundant individual NCPs compared to nucleosomes linked into arrays. However, TAZ2-mediated DNA binding enhances affinity of CZT for nucleosomal substrates, effectively sequestering CZT on initially bound NCPs by preventing dissociation and re-binding to new substrate NCPs, favouring activity of CZT on nucleosomes joined together as nucleosome arrays.

Moreover, CZT was markedly more active towards H3K27 and H2BK5 in nucleosome arrays compared to CZ (Figure 4C, D), consistent with TAZ2 specifically promoting acetylation of these physiologically important residues. This site-specific boost in activity suggests that TAZ2 DNA binding may not only function by simply increasing the residence time of CBP on chromatin, but might also facilitate interaction with the nucleosome in a conformation that supports efficient acetylation specifically of these physiologically important residues. In line with such a mechanism, CZT enzyme exhibited increased activity towards H3K27 and H2BK5 but reduced activity towards H3K9 and H4K5 compared to CZ (Figure 1D, E). These findings suggest that the TAZ2 domain promotes a specific mode of nucleosome engagement that favours H3K27 and H2BK5 acetylation at the expense of other potential substrate residues.

Lastly, to further confirm the role of TAZ2-mediated DNA binding in regulating CBP activity, we sought to identify mutations that would abrogate DNA binding. We identified three highly conserved, positively charged residues (R1769, K1832 and K1850 in mouse CBP) within TAZ2 that could play a role in DNA binding while not being expected to be important for overall structural integrity of the domain (Figure S2A). Indeed, a mutant TAZ2 domain (TAZ2^R1769E/K1832E/K1850E^, referred to as TAZ2^mut^) exhibited markedly diminished, although not completely abrogated, binding to both free DNA and NCPs (Figure 4E, S2D, S3C, D). Introduction of these mutations into the CZT construct decreased overall levels of acetylation towards all four core histone proteins, but most prominently towards histone H3 (Figure 4F). Moreover, when testing activity towards specific residues, acetylation of H3K27 and H2BK5 was most strongly compromised (Figure 4G, H). These results further support a DNA binding-dependent, TAZ2-mediated stabilization of CBP interaction with chromatin, promoting both overall histone acetylation activity as well as specificity towards H3K27.

### TAZ2 DNA binding promotes CBP specificity towards H3K27 in mouse ES cells

Having established that TAZ2-mediated DNA binding controls CBP activity and substrate specificity *in vitro*, we sought to test the role of TAZ2 DNA interactions in chromatin binding, histone acetylation, and gene expression *in vivo*. To this end, we used a catalytically inactive Cas9 (dCas9)-based epigenome editing system to target CBP fusion proteins to an endogenous gene promoter in mouse embryonic stem (ES) cells (Figure 5A) (Hilton et al., 2015). ES cells were co-transfected with four guide RNAs targeting the *Ascl1* promoter, which is inactive in ES cells and has been activated by dCas9-based epigenome editing previously (Black et al., 2016), and with HA-tagged dCas9 fused to CBP core, CZ, CZT^wt^, and CZT^mut^ constructs. In addition, fusions to catalytically inactive CZT (CZT^ci^, carrying a D1436Y point mutation) and to VP160 (Cheng et al., 2013) were used as negative and positive controls, respectively. Importantly, all dCas9 fusion constructs were expressed to similar levels (Figure 5B).

**Figure 5.**
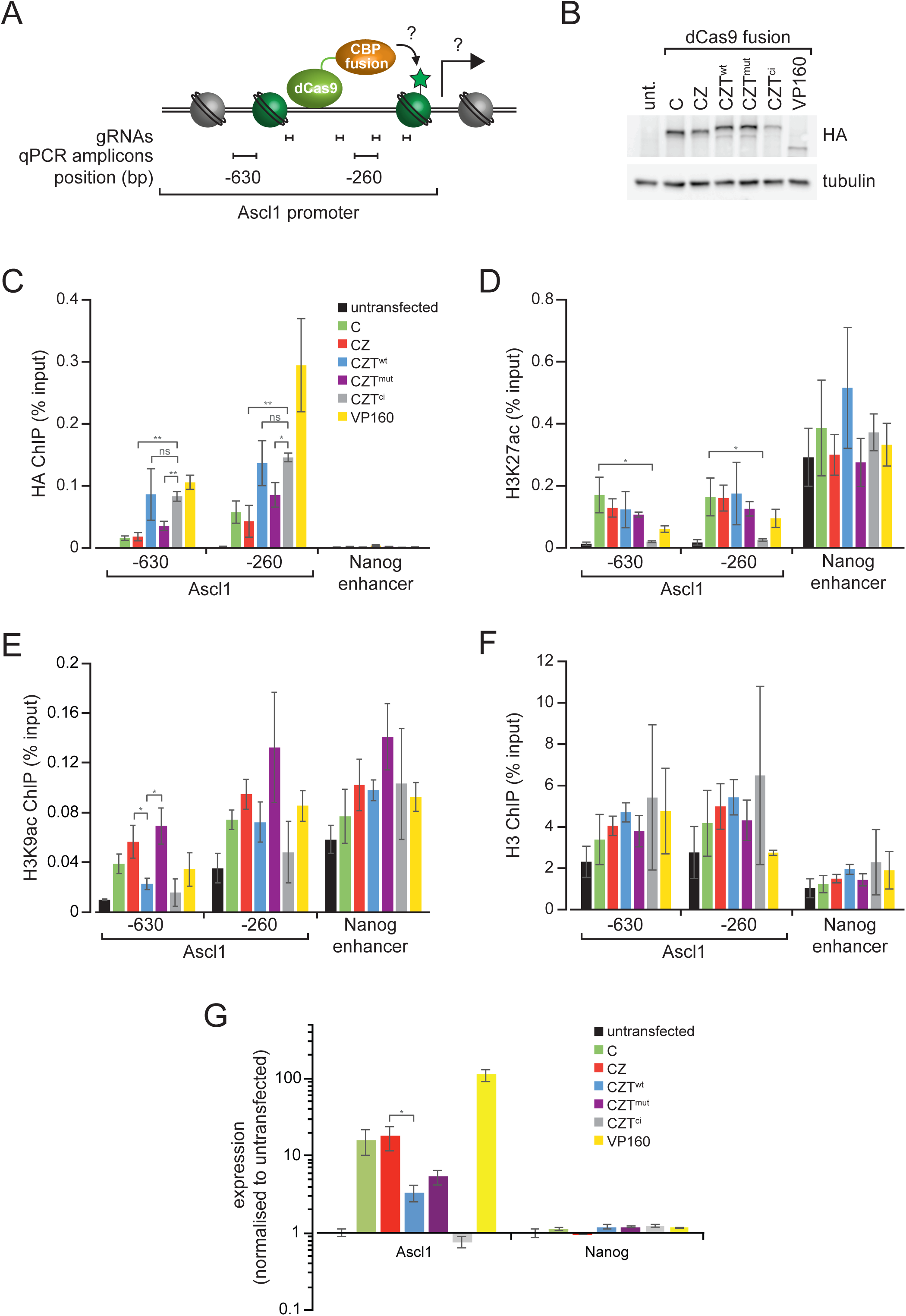
TAZ2 DNA binding promotes CBP specificity towards H3K27 in mouse ES cells. (**A**) Schematic showing targeting of dCas9-CBP fusion proteins to the mouse *Ascl1* promoter, with potential outcomes such as histone acetylation (green star) and gene transcription (arrow). Positions of bars indicate the locations of amplicons used for ChIP-qPCR relative to the transcription start site of *Ascl1*. ES cells were co-transfected with constructs expressing four *Ascl1* gRNAs and dCas9 fused to CBP core (C), CZ, CZT^wt^, CZT^mut^ (CZT with DNA-binding mutations), CZT^ci^ (CZT with catalytically inactivating mutations), or VP160 and analyzed after 48 h. (**B**) Whole cell extracts from transfected and untransfected (unt.) cells were analyzed by western blot for HA tag and tubulin as loading control. (**C–F**) ChIP qPCR analysis was carried out in transfected and untransfected cells, using antibodies against HA to measure binding of the dCas9 fusion protein (C), H3K27ac (D), H3K9ac (E) and histone H3 (F). qPCR was carried out at two regions within the *Ascl1* promoter and at the active Nanog enhancer as a control region. Data shown represents the mean and SEM of three independent transfections. *p<0.05, **p<0.01 (unpaired t-test). (**G**) Gene expression analysis was performed by RT-qPCR in transfected and untransfected cells, using primers for *Ascl1* and *Nanog* as a control gene. Gene expression is given relative to *Gapdh*, normalized to expression levels in untransfected cells, and shown on a logarithmic scale. Signal represents the mean and SEM of three independent transfections. *p<0.05 (unpaired t-test).

To test whether inclusion of the ZZ and TAZ2 domains affects binding of the dCas9-fusion proteins to the target site, chromatin immunoprecipitation (ChIP) was carried out. All dCas9 fusion proteins were robustly detected at the *Ascl1* locus but absent from the non-target *Nanog* enhancer (Figure 5C). However, whilst dCas9-CZ binding levels were unchanged compared to dCas9-CBP core, the TAZ2-containing dCas9-CZT^wt^ generated higher levels of chromatin binding (Figure 5C). Importantly, this increase in binding was lost for dCas9-CZT^mut^ but not dCas9-CZT^ci^. TAZ2-dependent enhanced association with chromatin in ES cells was also observed for CBP constructs lacking the dCas9 fusion partner (Figure S3E). These data indicate that the TAZ2 domain stabilizes CBP chromatin association in a manner that is dependent on its DNA binding activity but independent of CBP catalytic activity.

To test whether these differences in fusion protein binding result in differences in histone acetylation, ChIP experiments were carried out using antibodies against H3K27ac and H3K9ac (Figure 5D, E). While nucleosome occupancy remained unchanged for all constructs tested (Figure 5F), all constructs except dCas9-CZT^ci^ generated similar robust increases in H3K27ac at the *Ascl1* promoter, while acetylation at the non-target *Nanog* enhancer remained unchanged (Figure 5D). Importantly, H3K27ac generated by CBP fusion proteins was dependent on the catalytic activity of the enzyme, as H3K27ac levels in cells expressing dCas9-CZT^ci^ remained at background levels. By contrast, levels of H3K9ac generated by the dCas9 fusion proteins were more variable. Inclusion of the TAZ2 domain reduced levels of H3K9ac at the *Ascl1* promoter compared to dCas9-CBP core and dCas9-CZ (Figure 5E). Moreover, mutation of the TAZ2 domain in dCas9-CZT^mut^ led to increased H3K9ac to levels similar to those generated by dCas9-CBP fusions lacking TAZ2 (Figure 5E). Together, these results suggest that TAZ2 DNA binding facilitates activity towards H3K27 whilst attenuating activity towards non-target residues such as H3K9, thereby enhancing specificity of CBP *in vivo*. This is consistent with *in vitro* experiments demonstrating that the TAZ2 domain promotes acetylation of H3K27 while reducing the rate of reaction towards other potential substrates (Figure 1D, E).

Finally, to test whether TAZ2-mediated DNA binding influences transcriptional regulation, expression of *Ascl1* was assessed. While all dCas9 fusion proteins with the exception of dCas9-CZT^ci^ increased mRNA levels of *Ascl1*, dCas9-CZT^wt^ led to lower transcriptional activation than dCas9-CBP core and dCas9-CZ (Figure 5G), correlating with levels of H3K9ac generated (see Figure 5E). Diminished DNA binding in dCas9-CZT^mut^ resulted in approximately two-fold higher levels of gene expression (Figure 5G), mirroring the concomitant increase in H3K9ac (Figure 5E). These results suggest a model in which TAZ2 DNA binding enhances specificity of CBP towards H3K27, leading to reduced off-target acetylation and limiting the capacity of CBP to activate spurious transcription.

## Discussion

The acetyltransferases CBP/p300 are crucial for the regulation of gene expression in metazoans through placement of specific histone acetylation marks, most prominently H3K27ac. Here, we show that the TAZ2 domain of CBP is required for efficient acetylation of H3K27 in chromatin substrates *in vitro*. The TAZ2 domain binds DNA in a sequence-independent manner and promotes interaction between CBP and nucleosomes, thereby enhancing enzymatic activity and modulating substrate specificity of CBP. We further show that TAZ2 is required to stabilize CBP binding to chromatin *in vivo* and to facilitate activity and specificity towards H3K27. Together, these results uncover a novel mechanism by which CBP interacts with chromatin and elucidate a new mode of regulation for this important histone modifier.

Previous work investigating CBP and p300 enzymatic activity suggested an inherent promiscuity towards histones *in vitro* that is not observed *in vivo* (An and Roeder, 2003; Ogryzko et al., 1996; Weinert et al., 2018). Recent work found that the ZZ domain specifically promotes acetylation of H3K27 in nucleosomes through interaction with the N-terminal tail of histone H3 (Zhang et al., 2018). While we similarly observed an increase in H3K27 acetylation mediated by the ZZ domain, we found ZZ to increase acetylation towards multiple lysines in nucleosomes, not only towards H3K27 (Figures 1, 4). This discrepancy might be caused by differences in experimental conditions. While we performed HAT assays with a two-fold molar excess of nucleosome substrate over enzyme, previous work (Zhang et al., 2018) used a three-fold molar excess of enzyme. Under conditions of excess enzyme, acetylation of residues such as H3K9 could have been saturated and thus appeared unchanged, whereas less readily modified residues such as H3K27 exhibited increased acetylation via the ZZ domain. By contrast, we found the TAZ2 domain to both robustly enhance activity of CBP and increase specificity for H3K27.

The ability of the TAZ2 domain to bind DNA and change the substrate preferences of CBP to favour acetylation of H3K27 appears to further explain mechanistically how assembly of histones into chromatin modulates CBP activity and substrate selection (An and Roeder, 2003; Kraus et al., 1999). Moreover, the key role played by the TAZ2 domain in CBP function may also rationalize recent findings that missense mutations within TAZ2 expected to affect its structural integrity are associated with the rare developmental disorder Menke-Hennekam syndrome (MKHK) (Banka et al., 2019; Menke et al., 2018; Menke et al., 2016). Importantly, MKHK patients carrying mutations in key zinc-coordinating residues that are likely to destabilize the TAZ2 domain display a clinical phenotype that is distinct from patients with Rubinstein-Taybi syndrome (RSTS), which is caused by loss of CBP function (Hennekam, 2006; Menke et al., 2016). In contrast, MKHK clinical features resemble those found in patients with CBP duplication, suggesting that MKHK is the result of CBP gain of function mutations (Menke et al., 2018). Our observations that deletion of the TAZ2 domain is not associated with abrogation of enzymatic activity but rather with modulation of CBP substrate specificity, leading to promiscuous acetylation activity of CBP, are consistent with a gain of function phenotype. The resulting relaxed substrate specificity may result in aberrant gene expression (see below, Figure 6), potentially explaining some of the phenotypes associated with MKHK.

**Figure 6.**
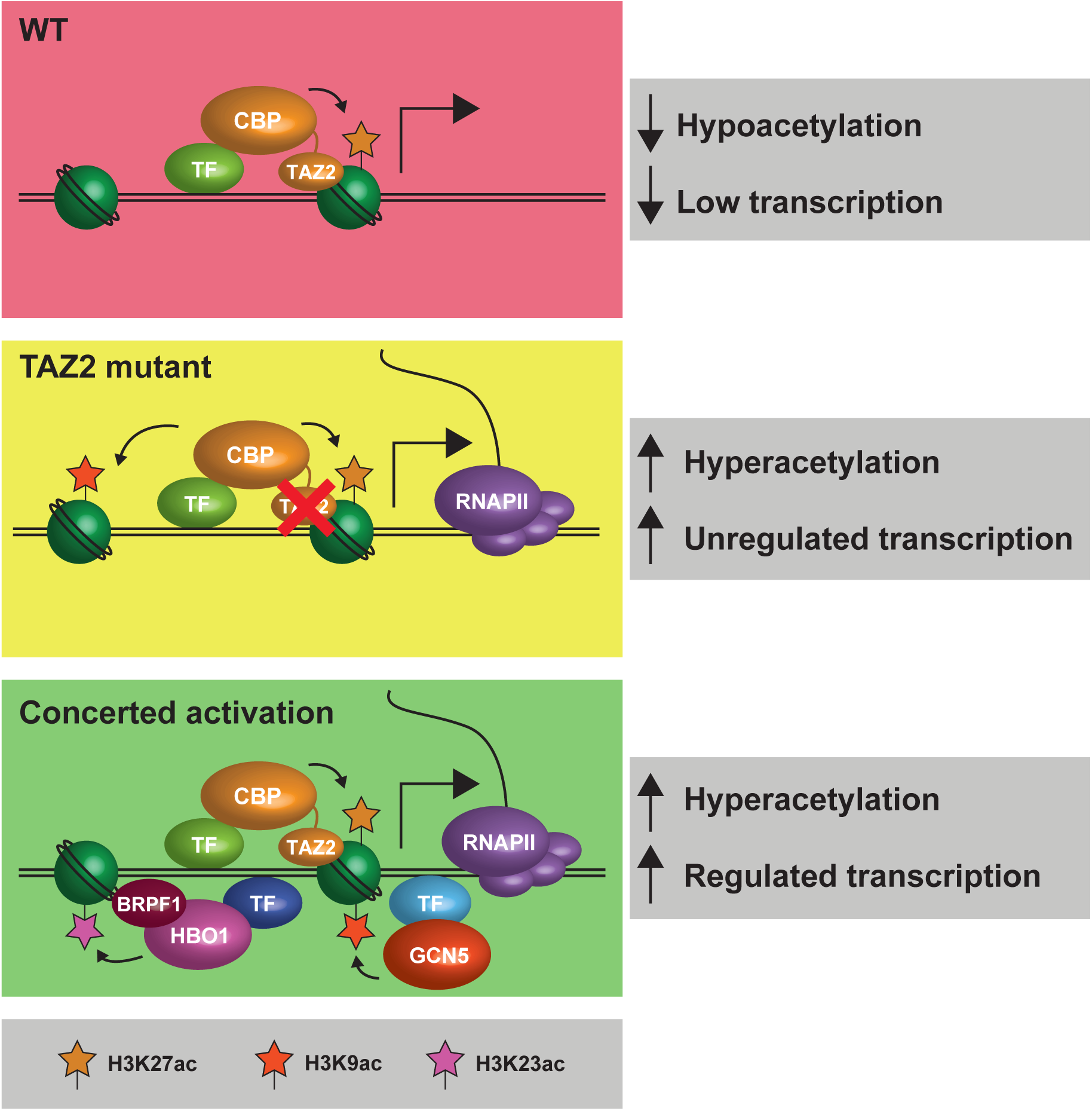
A model for CBP-mediated histone acetylation in transcriptional regulation. A model of histone acetylation at a gene regulatory element. When the activity of CBP is restrained by the TAZ2 domain, CBP-mediated H3K27ac by itself is unable to support high levels of transcription (top). Mutation of the TAZ2 domain leads to more promiscuous acetylation by CBP, facilitating unregulated transcriptional activation (middle). Regulated transcriptional activation requires the concerted activity of multiple HATs specifically acetylating multiple histone residues (bottom).

More generally, these results also illustrate the importance of sequence-independent DNA interactions for multivalent engagement of nucleosomes by chromatin-modifying proteins (Ruthenburg et al., 2007). Similarly to the function revealed here for the TAZ2 domain in CBP, MOZ features a DNA-binding zinc finger within its HAT domain (Holbert et al., 2007). Moreover, the double PHD finger of the closely related HAT MORF also binds DNA in a sequence-independent fashion, crucially contributing to interactions between MORF and nucleosomes (Klein et al., 2019). In CBP and other HATs, such sequence-independent DNA binding could function to stabilize the enzyme on chromatin regardless of the underlying DNA sequence, cooperating with transcription factor-based recruitment to facilitate longer residence time at target sites and to increase productive interactions with nucleosome substrates.

In addition to enhancing overall enzymatic activity, we found TAZ2 DNA binding to modulate CBP substrate selection by specifically favouring acetylation of H3K27, a function which cannot solely be accounted for by increased residence time. We reason that the TAZ2 domain may therefore function analogously to the chromobarrel domain of MOF, whose DNA binding activity is specifically required to trigger acetylation of H4K16 (Conrad et al., 2012). Further mechanistic parallels can be drawn with HBO1, which forms mutually exclusive complexes with the homologous partner proteins BRPF1 and JADE1 (Lalonde et al., 2013). The zinc knuckle and second PHD finger within the PHD-Zn knuckle-PHD (PZP) region of BRPF1 binds DNA in a sequence-independent manner, whilst key residues mediating this interaction are not conserved in JADE1 (Klein et al., 2016). While BRPF1 is required for HBO1 activity towards histone H3 specifically in the context of chromatin, interaction with JADE1 leads to a switch in HBO1 acetylation specificity towards H4 (Lalonde et al., 2013). This subunit-dependent substrate switch suggests that HBO1 requires sequence-independent DNA binding for specificity towards its major targets, H3K14 and H3K23, similar to the requirement of TAZ2-mediated DNA binding for H3K27 acetylation by CBP. Interestingly, while the zinc knuckle and second PHD finger of the PZP region in BRPF1 bind DNA, the first PHD finger immediately upstream interacts with the unmodified N-terminal tail of histone H3 (Klein et al., 2016), a further mechanistic parallel with the tandem arrangement of ZZ and TAZ2 domains in CBP. This suggests that DNA binding, augmented by additional histone tail interactions, may therefore be a commonly employed mechanism to generate substrate specificity for histone-modifying enzymes.

DNA binding appears to be particularly important for acetylation of residues that lie in close proximity to the gyres of DNA such as H3K27, H3K23, and H4K16 (Davey et al., 2002; Luger et al., 1997), where proximity with nucleosomal DNA could hinder access of HAT domains to their substrates. In addition to increasing residency time on nucleosomes, binding to nucleosomal DNA by TAZ2 may therefore also orient the enzyme in a topology that favours interaction with, and subsequent acetylation of, H3K27. Beyond HATs, such a model would also be consistent with recent work showing that interactions with nucleosomal DNA are required for positioning of Polycomb repressive complex 2 (PRC2) for efficient methylation of H3K27 (Finogenova et al., 2020).

Finally, the dCas9-based targeting experiments presented here add to our understanding of the role of H3K27ac in transcriptional activation. Consistent with recent observations that H3K27ac is not uniquely important for maintaining gene expression (Zhang et al., 2020), we found that the presence of H3K27ac alone is not sufficient to generate high levels of transcription, but requires co-occurrence of at least H3K27ac and H3K9ac (Figure 5). These observations support a model in which multiple histone acetylation marks, including H3K27ac, function in concert to maintain full and robust transcriptional activation of target genes, while enabling complex regulation and integration of multiple inputs via concerted action of multiple HATs, specific acetylation marks, their cognate readers and direct impact on chromatin structure (Figure 6). Specificity of individual HATs for individual histone lysine residues would be pivotal for such a system. By focusing CBP specificity towards a narrow set of histone substrates, the TAZ2 domain could therefore play a central role in maintaining such transcriptional robustness. Moreover, sequence-independent DNA binding by HATs could represent a common mechanism by which substrate specificity is achieved, allowing HATs to cooperate to maintain robustness of the gene expression programme.

## Acknowledgments

We thank Shelley Berger for the pFastBac CBP plasmid. The pAC94-pmax-dCas9VP160-2A-puro plasmid (Addgene 48226) was a kind gift from Rudolf Jaenisch and the pmU6-gRNA plasmid (Addgene 53187) was a kind gift from Charles Gersbach. We are grateful to Adrian Bird and Robin Allshire for critical reading of the manuscript. We thank Adrian Bird, Atlanta Cook, Ken Sawin, Marcus Wilson, and Anca Farcas for helpful discussions and insightful comments on this work. We also thank all members of the Voigt for helpful discussions.

## Author contributions

T.W.S., Conceptualization, Methodology, Formal analysis, Investigation, Data curation, Writing—original draft preparation, Writing—review & editing, Visualization, Project administration. V.M., K.W., E.B., Resources. P.V., Conceptualization, Validation, Resources, Writing—original draft preparation, Writing—review & editing, Visualization, Supervision, Project administration, Funding acquisition.

## Figure Supplement Legends

**Figure S1.**
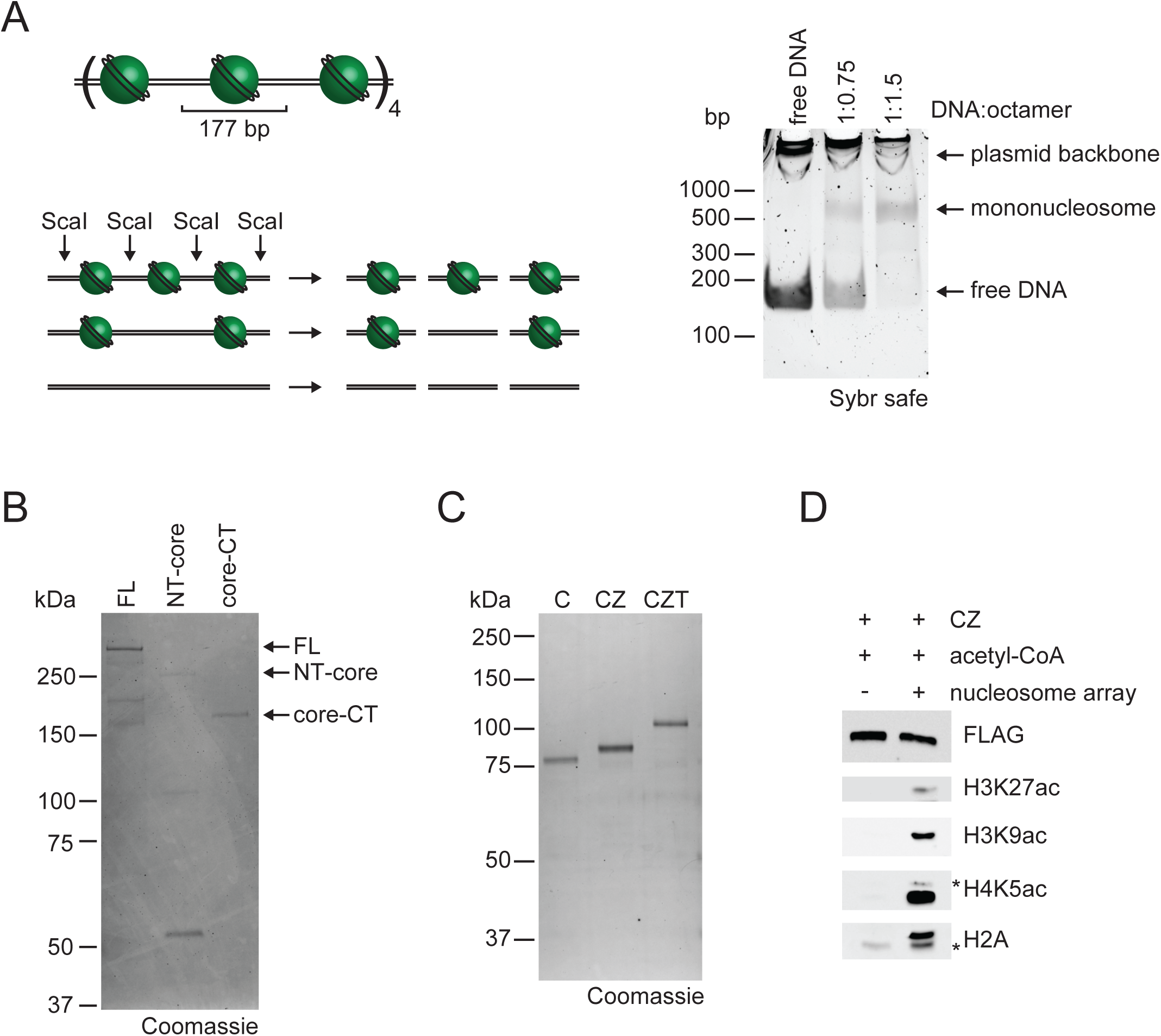
Assembly of nucleosome arrays and expression of CBP proteins. (**A**) Nucleosome arrays comprise 12 repeats of a 177-bp sequence containing the 601 nucleosome positioning sequence (left, top). Nucleosome arrays were reconstituted using different DNA:histone octamer mass ratios and analyzed by digestion with the restriction endonuclease ScaI to release either free DNA from unoccupied positioning sequences or mononucleosomes from nucleosome-occupied sequences (shown schematically on left). Digested arrays were analysed by native PAGE (right), showing that a 1:1.5 ratio of DNA:histone octamer led to full occupancy of the nucleosome array, as the free DNA band was absent. (**B**) Purified CBP full-length (FL), NT-core and CT-core proteins were analyzed by SDS-PAGE and Coomassie staining. (**C**) Purified CBP core, core-ZZ (CZ), and core-ZZ-TAZ2 (CZT) proteins were analyzed by SDS-PAGE and stained with Coomassie. (**D**) Western blot analysis of HAT assays using CZ construct shows that non-specific bands (marked by asterisks) observed with H4K5ac and H2A antibodies are primarily present only in the presence of acetyl-CoA and therefore likely correspond to non-specific signal from acetylated histones.

**Figure S2.**
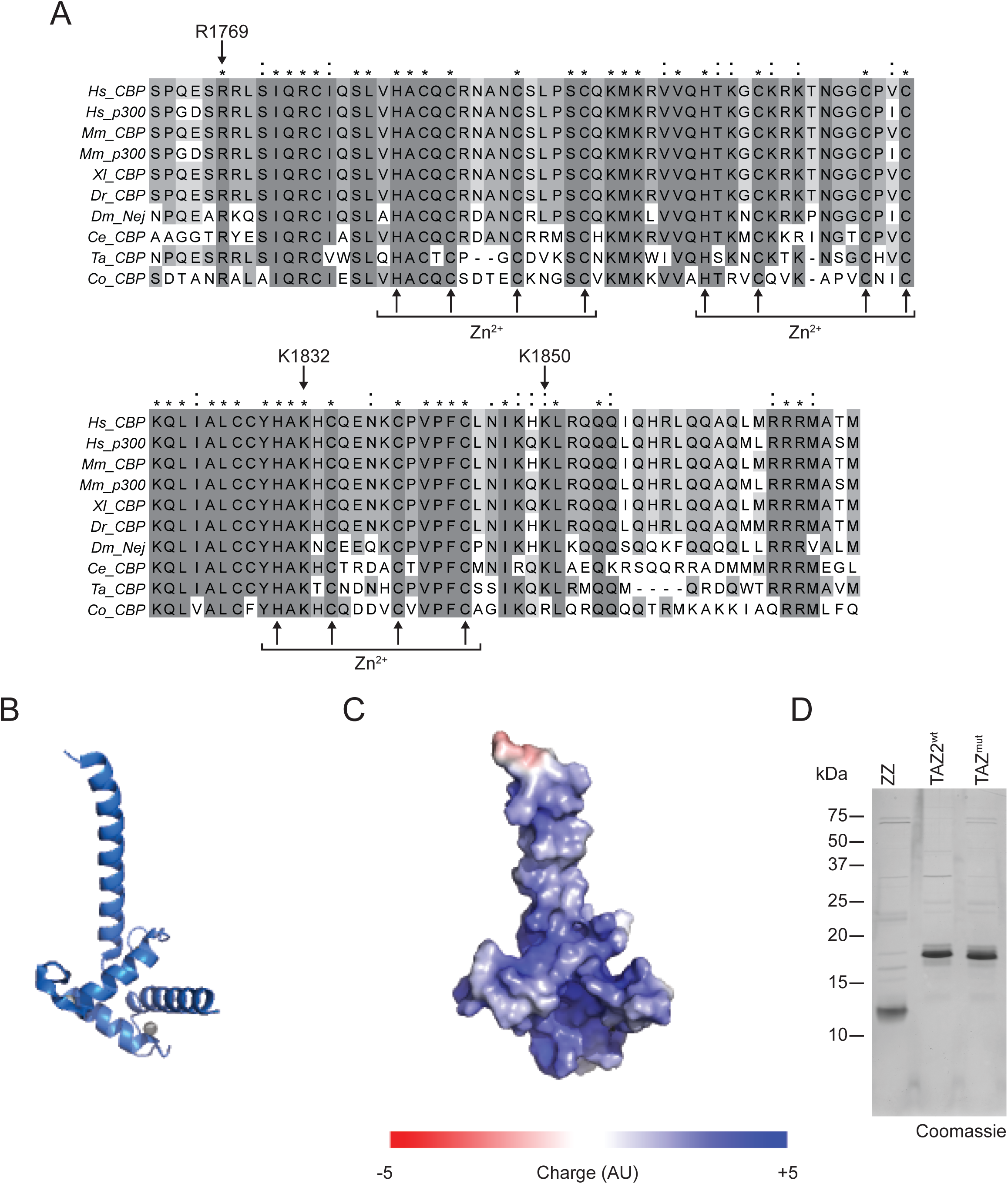
The TAZ2 domain is a highly conserved basic domain. (**A**) Alignment of TAZ2 domains in CBP and p300 proteins in humans (*Hs*), mice (*Mm*), *Xenopus laevis* (Xl), *Danio rerio* (*Dr*), *Drosophila melanogaster* (*Dm*), *Caenorhabditis elegans* (*Ce*), *Trichoplax adhaerens* (*Ta*), and *Capsaspora owczarzaki* (*Ca*). Greyscale shading represents level of conservation, asterisks (*) indicate identical residues and colons (:) represent similar residues. Arrows underneath and above the sequences indicate zinc-coordinating residues and residues that were mutated to abrogate DNA binding, respectively. Protein sequences were obtained from UniProtKB, multiple sequence alignment was carried out with Clustal Omega and alignments were visualised using JalView. (**B**) Cartoon representation of the crystal structure of human p300 TAZ2 domain (PDB ID: 3IO2) showing protein backbone in blue and coordinated zinc ions as grey spheres. (**C**) Surface representation of the structure in (B) with colour-coded surface charge calculated using PyMol APBS. (**D**) ZZ, TAZ2^wt^, and TAZ2^mut^ proteins were expressed and purified from bacteria, analysed by SDS-PAGE and stained with Coomassie.

**Figure S3.**
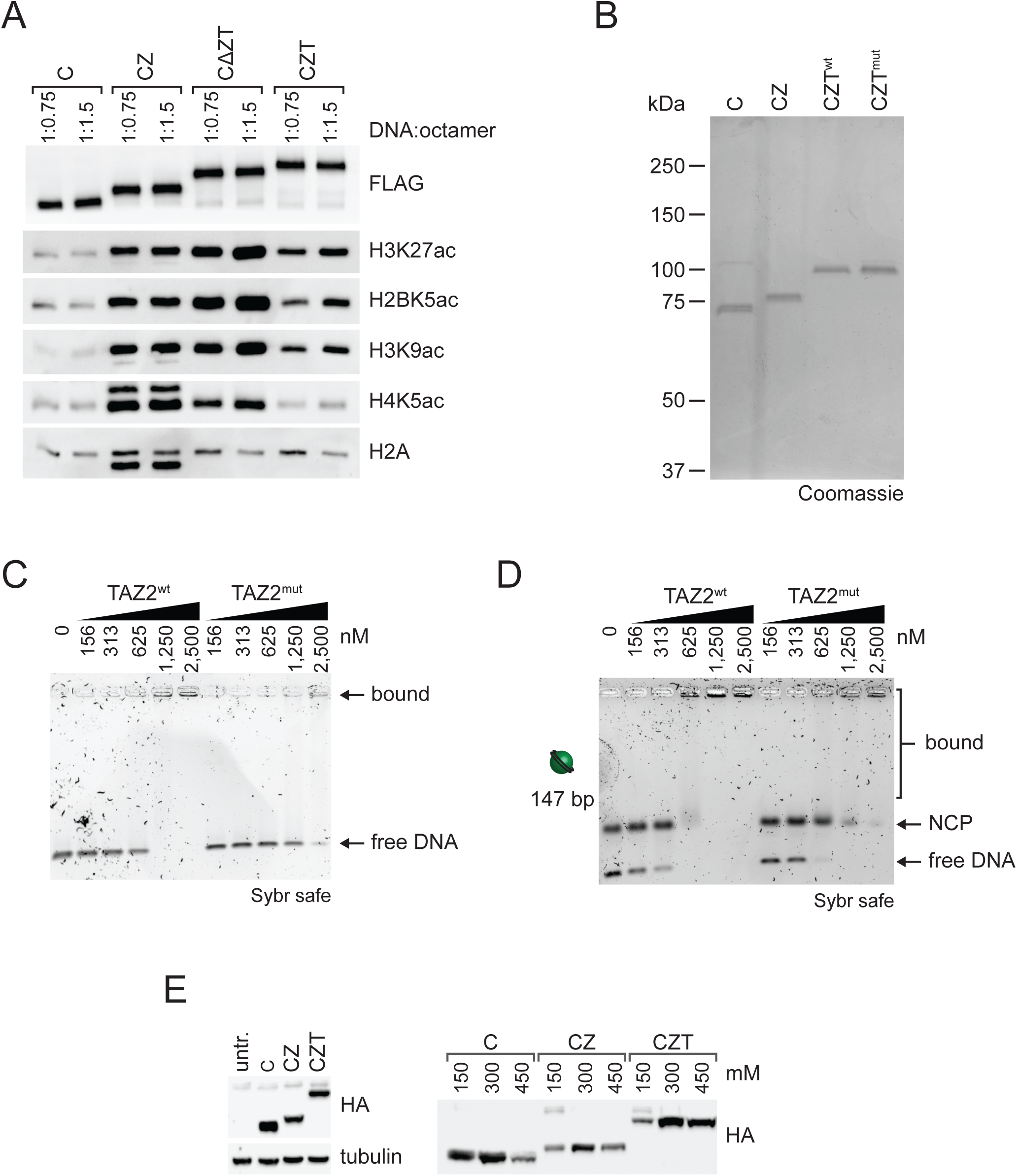
TAZ2 DNA binding affects interactions with chromatin. (**A**) HAT assays were carried out with nucleosome arrays reconstituted with different ratios of DNA:histone octamer, yielding different numbers of nucleosomes per DNA molecule. Reactions were performed with 150 nM of nucleosome (by histone octamer) and 150 nM CBP core (C), core-ZZ (CZ), core-ΔZZ-TAZ2 (CΔZT), and core-ZZ-TAZ2 (CZT) enzymes. The reactions were then analysed by western blot using antibodies against FLAG, H3K27ac, H2BK5ac, H3K9ac, H4K5ac, and total H2A. Data shown is representative of three independent experiments. (**B**) Purified CBP core, CZ, CZT^wt^, and CZT^mut^ enzymes were analysed by SDS-PAGE and stained with Coomassie. (**C, D**) EMSA analysis of binding of TAZ2^wt^ or TAZ2^mut^ domains to either free 147-bp 601 DNA (C) or NCPs (D). Experiments shown are representative of three independent EMSAs. **(E)** Left panel: Whole cell extracts from ES cells transfected with constructs expressing CBP core (C), CZ, and CZT were analysed by western blot using antibodies against HA and tubulin. Right panel: Nuclei isolated from transfected cells were sequentially eluted with nuclear extract buffer containing 150, 300, and 450 mM NaCl to determine strength of chromatin binding, with elution in higher salt indicating stronger association with chromatin. Nuclear extracts were analysed by western blot using anti-HA antibody.

## Materials and Methods

### DNA constructs for protein expression

Full-length N-terminally His-FLAG-tagged CBP in pFastBac HTA (Bose et al., 2017) was a kind gift from Shelley Berger (University of Pennsylvania). Sequencing of this construct revealed a point mutation that generates a P695L mutation in an unstructured region of CBP, which was carried forward in the CBP core-NT construct and is not expected to impact CBP catalytic activity. CBP truncations were generated by PCR amplification with suitable primers, and the ZZ domain deletion construct was generated by overlap-extension PCR. All CBP constructs were cloned into a modified pFastBac plasmid to yield in-frame fusion with a 5’ sequence encoding a 6xHis tag, TEV protease cleavage site, and a 1xFLAG tag. The TAZ2^mut^ sequence was synthesised by IDT and cloned into the pFastBac vector using an internal XhoI site in the HAT domain of CBP. For bacterial expression, isolated ZZ and TAZ2 domains were cloned into a modified pET22 vector in frame with a 5’ sequence encoding an N-terminal 6xHis tag and TEV cleavage site.

### Protein expression and purification from Sf9 cells

Sf9 cells were maintained in suspension cultures in serum-and antibiotic-free HyClone CCM3 media (GE Life Sciences) at 27°C and 125 rpm. Recombinant baculoviruses were generated by transfection of Sf9 cells with bacmid generated through transformation into EmBacY cells, using X-tremeGENE HP transfection reagent (Roche). After 3–4 rounds of virus amplification, Sf9 cells were infected for 48 h at previously optimized virus titers for protein expression.

Insect cell pellets from 500 ml of culture were collected by centrifugation and resuspended in 30 ml lysis buffer (20 mM Tris-Cl pH 8.0, 350 mM NaCl, 10% glycerol, 10 μM ZnCl_2_, 0.1% NP40, 0.5 mM PMSF, 1 mM DTT) and lysed by sonication. The lysate was cleared by centrifugation at 40,000 x *g* for 30 min at 4°C and the supernatant filtered through a syringe-driven 0.45-μm PVDF membrane (Millipore). To prepare the beads, 100 μl of packed FLAG M2 affinity resin (Sigma) was washed twice with BC100 (20 mM Tris-Cl pH 8.0, 100 mM NaCl, 10% glycerol, 1 mM DTT), once with 0.1 M glycine pH 2.5, twice with 1 M Tris-Cl pH 8 and twice with lysis buffer, with the resin collected after each wash step by centrifugation at 800 x g for 2 min. The resin was then added to the cleared lysate and incubated for 2 h at 4°C with end-over-end rotation. The bound resin was washed three times with BC350 wash buffer (20 mM Tris-Cl pH 8.0, 350 mM NaCl, 10% glycerol, 10 μM ZnCl_2_, 0.5 mM PMSF, 1 mM DTT) and once with BC100. Bound protein was eluted three times for 30 min by incubation with 0.3 mg/ml 3xFLAG peptide (Sigma) in BC100 at 4°C with end-over-end rotation, snap frozen and stored at -80°C.

### Protein expression and purification from bacteria

ZZ and TAZ2 domain were expressed in BL21 (DE3) *E. coli*. Cultures were grown at 37°C until OD_600_ was approximately 0.6, when protein expression was induced for 3 h at 37°C by addition of 0.5 mM IPTG and 20 μM ZnCl_2_. The bacteria were harvested by centrifugation at 6,000 x *g* for 15 min at 4°C, washed once with 1x PBS, flash frozen, and stored at -80°C.

Bacterial pellets were resuspended in 16 ml lysis buffer (20 mM Tris-Cl pH 8, 500 mM NaCl, 0.1% NP40, 0.5 mM PMSF) and lysed by sonication. The lysate was cleared by centrifugation at 23,000 x *g* for 30 min at 4°C. The 6xHis tagged protein was purified by IMAC affinity purification using 0.5 ml of packed Sepharose 6 Fast Flow Ni-NTA Resin (GE Healthcare) pre-washed with lysis buffer. The binding reaction was allowed to proceed for 1 h at 4°C with end-over-end rotation. The beads were then pelleted by centrifugation at 800 x *g* for 2 min at 4°C and the unbound flowthrough fraction was removed. The beads were resuspended in 20 column volumes (CV) of low salt wash buffer (50 mM NaH_2_PO_4_ pH 8, 300 mM NaCl, 20 mM imidazole, 0.1 mM PMSF), added to a 10-ml poly-prep chromatography column (Bio-Rad) and the wash buffer allowed to flow through the column. The column was then washed with 20 CV high salt wash buffer (50 mM NaH_2_PO_4_ pH 8, 1 M NaCl, 20 mM imidazole, 0.1 mM PMSF), and subsequently equilibrated into low salt with 10 CV low salt wash buffer. Bound proteins were eluted 5-10 times with 1 CV elution buffer (50 mM NaH_2_PO_4_ pH 8, 300 mM NaCl, 250 mM imidazole). Protein concentration and purity in the elution fractions was determined by SDS-PAGE followed by Coomassie staining with Instant Blue (Expedeon). The most concentrated and pure fractions were pooled and dialysed against BC100 buffer (20 mM HEPES pH 8, 100 mM KCl, 10% glycerol, 0.5 mM DTT).

The dialysed protein was cleared of precipitate by centrifugation at 13,000 x *g* for 10 min at 4°C. The purified protein was snap frozen and stored at -80°C.

### Nucleosome reconstitution

Nucleosome arrays for HAT assays were reconstituted onto a plasmid called p177-601 containing 12 copies of the 601 nucleosome positioning sequence (Lowary and Widom, 1998) each separated by a 30-bp linker. For EMSAs using nucleosome arrays, nucleosomes were reconstituted onto linear 12x 601 array, which was generated by excision of the array from p177-601 by digestion with EcoRV. To reconstitute NCPs, 147 bp and 209 bp 601 DNA was amplified by PCR using a biotinylated forward primer, and PCR products were purified using the EZNA Gel Extraction kit (Omega BioTek).

Histone octamers were assembled as described previously (Voigt et al., 2012). In short, human histones H2A and H2B and *Xenopus laevis* histones H3 and H4 were expressed in *E. coli*, purified from inclusion bodies and solubilised in unfolding buffer (20 mM Tris-Cl pH 7.5, 7 M guanidine HCl, 10 mM DTT). After dialysis into urea buffer (10 mM Tris-Cl pH 8, 100 mM NaCl, 7 M urea, 1 mM EDTA, 5 mM β-mercaptoethanol), histones were further purified by tandem ion exchange chromatography using HiTrap Q and Hi Trap SP anion and cation exchange columns (GE Life Sciences). Histone-containing fractions were pooled, dialysed against 3 mM β-mercaptoethanol in water, and lyophilised for storage at -80°C. To reconstitute histone octamers, histone proteins were resuspended in unfolding buffer and mixed in equal amounts but with an 20% excess of H2A and H2B to ensure complete octamer formation. After dialysis against refolding buffer (10 mM Tris-Cl pH 8, 2 M NaCl, 1 mM EDTA, 5 mM β-mercaptoethanol), refolded octamers were purified using an S200 gel filtration column (GE Life Sciences).

To reconstitute nucleosomes, DNA template and histone octamer were mixed in refolding buffer at a DNA:histone octamer mass ratio of 1:2 for NCPs and 1:1.5 for nucleosome arrays, unless otherwise stated. The assembly reactions were then subjected to gradient dialysis with TE buffer to a final NaCl concentration of 400 mM, followed by step dialysis against TE buffer. Proper reconstitution of NCPs was checked by analysis on a 1% agarose gel in TBE buffer, and reconstitution of nucleosome arrays was verified by digestion with the restriction enzyme ScaI at 37°C for 1 h followed by analysis on a 6% native PAGE gel.

### Histone acetyltransferase (HAT) assays

Enzyme and substrate were mixed on ice in 1 x HAT buffer (50 mM Tris-Cl pH 7.5, 10% glycerol, 4 mM DTT) to a final concentration of 75 nM enzyme and 150 nM substrate (approximately 425 ng of histone protein) in a final reaction volume of 25 µl. For radiolabel-based assays, reactions were started by addition of [^3^H]-acetyl coenzyme A (acetyl-CoA) (Hartmann Analytic) to a final activity of 1.5 µCi (corresponding to 7.8 µM) per reaction. For unlabelled assays, unlabelled acetyl-CoA (Sigma) was added to a final concentration of 100 µM. Reactions were allowed to proceed for 20 min (unless indicated otherwise) at 30°C in a ThermoMixer (Eppendorf) shaking at 550 rpm and were stopped by addition of 1xSDS loading buffer and boiling for 5 min at 95°C.

Quenched reactions were then separated by SDS-PAGE on 18% and 8% gels to analyse histone proteins and CBP constructs, respectively. For [^3^H]-labelled HAT assays, proteins were then transferred to a PVDF membrane using the TurboTransfer system (Bio-Rad). The membrane was stained with Coomassie, destained, and air dried. Once dry, the membrane was exposed to film (Carestream Kodak BioMax MS film, Sigma) through an intensifying screen (BioMax Transcreen, Sigma) at -80°C for 1–5 days before the autoradiograph was developed. For non-radioactive HAT assays, proteins were transferred to nitrocellulose membranes and analysed by western blot with the antibodies listed in Table 1. Chemiluminescence was detected with a ChemiDoc Touch system (Bio-Rad) and signals quantified with the Volume tool in Image Lab software (Bio-Rad).

**Table 1.**
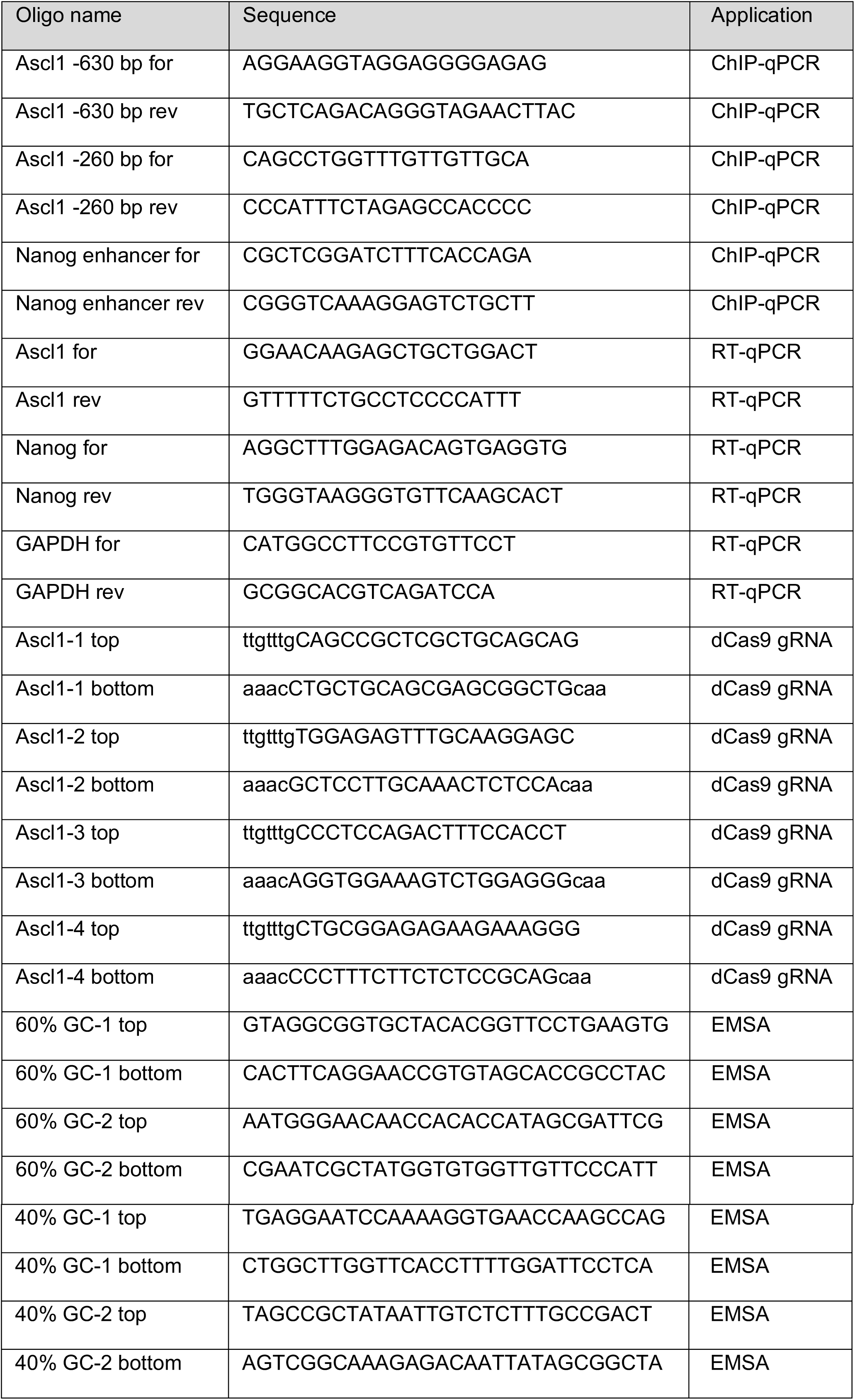
Oligonucleotides used in this study.

### Electrophoretic mobility shift assays (EMSAs)

To generate 29 bp dsDNA probes for EMSA, complementary 29-nt oligonucleotides (see Table 1 for sequences) were each reconstituted in H_2_O at 100 µM concentration and 25 µl of each oligo were mixed for annealing. The oligos were annealed in a thermal cycler by incubating at 37°C for 30 min, boiling at 95°C for 5 min, and then reducing the temperature in 5°C intervals, holding each temperature for 1 min, to a final temperature of 25°C.The duplex DNAs were then diluted for use in binding reactions.

Binding reactions for EMSA experiments were set up in a final volume of 10 µl in 1x EMSA buffer (20 mM HEPES pH 8, 150 mM KCl, 7.5% glycerol, 0.5 mM DTT). DNA or nucleosomes were added to a final concentration of 35 nM of DNA or mononucleosome equivalent. Binding reactions were started by the addition of protein, gently mixed, and incubated on ice for 30 min.

Binding reactions were analysed by agarose gel electrophoresis in 0.5x TBE, using 1.2% agarose for 29 bp dsDNA probes, 1% agarose for free 147 bp 601 DNA, 1% agarose for 147 or 209 bp nucleosome core particles (NCPs), and 0.5% agarose for 12x nucleosome arrays. 8.5 µl of each sample was loaded and electrophoresis carried out at 100 V (8.3 V/cm) for 20 min for 1.2% gels, 30 min for 1% gels and 40 min for 0.5% gels, in 0.5x TBE buffer at room temperature. The gel was then stained in 1x Sybr safe for 15 min and imaged using the ChemiDoc Touch system. EMSAs were analyzed by quantifiying the unbound band using the Volume tool in Image Lab software. Dissociation constants were calculated by Hill function fitting with GraphPad Prism.

### DNA pulldown

For DNA pulldown experiments with ZZ or TAZ2 domains, 20 µl slurry of Streptavidin M-280 Dynabeads (ThermoFisher) were blocked twice with 0.5% bovine serum albumin (BSA) in 1x PBS and then washed twice with TEN buffer (10 mM Tris-Cl pH 8, 1 mM EDTA, 1 M NaCl). To bind DNA, the beads were resuspended in 1 ml of TEN buffer, 1 µg of biotinylated 147 bp 601 DNA was added to the beads (or no DNA was added for control pulldowns), and incubated for 1 h at 4°C with end-over-end rotation. After washing the DNA-bound beads once with TEN buffer and twice with binding buffer (50 mM Tris-Cl pH 8, 50 mM NaCl, 0.05% NP40, 0.5 µg/ml BSA, 0.5 mM DTT), 1 µg of purified protein was added in 1 ml of binding buffer and incubated with rotation for 2-3 h at 4°C. To remove unbound protein, the beads were washed three times for 10 min with 1 ml of wash buffer (50 mM Tris-Cl pH 8, 300 mM NaCl, 0.05% NP40, 0.5 µg/ml BSA, 0.5 mM DTT), and the remaining bound protein was eluted by boiling for 5 min in 60 µl of 1x SDS-PAGE loading buffer. Binding was then analysed by separation of 10% of input protein and 50% of pulldown material on an 18% SDS-PAGE gel followed by western blot using anti-His antibody.

### Mammalian cell culture

E14 ES cells were grown at 37°C in 5% CO_2_ on gelatin-coated plates in media comprising high glucose DMEM (Gibco), 15% foetal bovine serum (Gibco), 100 µg/ml penicillin/streptomycin (Gibco), 2 mM L-glutamine (Gibco), 1x non-essential amino acids (Gibco), 50 µM β-mercaptoethanol (Sigma) and 10 ng/ml heterologously expressed home-made leukaemia inhibitory factor (LIF).

### dCas9-based genomic targeting

CBP sequences were cloned into the pAC94-pmax-dCas9VP160-2A-puro vector (Addgene: 48226, a kind gift from Rudolf Jaenisch) described by (Cheng et al., 2013) by excision of the VP160 coding sequence and replacing it with the CBP sequences. The four guide sequences for the *Ascl1* locus (see Table 1) were taken from (Black et al., 2016) and cloned into pmU6-gRNA (Addgene: 53187, a kind gift from Charles Gersbach) (Kabadi et al., 2014) for expression as single individual guide RNAs (sgRNAs). To express the dCas9 fusion proteins and sgRNAs, ES cells were transiently transfected with 15 µg of pAC94 plasmid DNA and 1.25 µg each of the four different pmU6 plasmids using Lipofectamine 3000 and OptiMEM (Gibco). Cells were seeded on a 15-cm plate 24 h before transfection and grown in ES media without penicillin/streptomycin. The morning after transfection, the media was replaced with complete ES media and cells were cultured for a further 24 h. 48 h after transfection, cells were harvested by trypsinisation, washed once with 1x PBS and divided into pellets for preparation of whole cell extracts, chromatin, and RNA.

### Whole cell extract

To prepare whole cell protein extracts from ES cells, pellets were resuspended in RIPA buffer (50 mM Tris-Cl pH 8.0, 150 mM NaCl, 10% glycerol, 1% NP40, 1 mM DTT, 0.5 mM PMSF). After lysis on ice for 30 min, cellular debris was pelleted by centrifugation at 15,000 x *g* for 20 min at 4°C. The supernatant was taken as whole cell extract.

### Chromatin immunoprecipitation (ChIP)

To prepare chromatin, 1x 10^7^ ES cells were resuspended in 10 ml of 1x PBS and crosslinked by addition of 16% methanol-free formaldehyde (Pierce) to a final concentration of 1% and incubated for 10 min at room temperature. The crosslinking reactions were quenched by addition of glycine to a final concentration of 125 mM and the cells were pelleted by centrifugation at 800 x *g* for 4 min. The crosslinked cells were lysed in 10 ml LB1 (50 mM HEPES pH 8, 140 mM NaCl, 1 mM EDTA, 10% glycerol, 0.5% NP40, 0.25% Triton X-100, 0.5 mM PMSF) for 10 min at 4°C with end-over-end rotation. The released nuclei were recovered by centrifugation at 800 x *g* for 4 min and were subsequently washed in 10 ml of LB2 (10 mM Tris-Cl pH 8, 200 mM NaCl, 1 mM EDTA, 0.5 mM EGTA, 0.1 mM PMSF) for 10 min at 4°C with end-over-end rotation. After centrifugation at 800 x *g* for 4 min, nuclei were resuspended in 1 ml of LB3 (10 mM Tris-Cl pH 8, 200 mM NaCl, 1 mM EDTA, 0.5 mM EGTA, 0.1% sodium deoxycholate, 0.5% N-lauroylsarcosine, 0.1 mM PMSF) and transferred to a 15-ml hard plastic polystyrene falcon tube (BD Falcon) for sonication. Chromatin was fragmented by sonication using a Bioruptor (Diagenode) on the high power setting for 30 min with pulses of 30 s on/30 s off, giving a total sonication time of 15 min and average fragment sizes of 200-300 bp. Following sonication, 10% Triton X-100 dissolved in LB3 was added to a final concentration of 1%. The chromatin was cleared by centrifugation at 15,000 x *g* for 10 min at 4°C and the supernatant was taken as the chromatin extract. Chromatin was then either used immediately for ChIP or aliquoted and stored at -80°C.

To immunoprecipitate chromatin, antibodies were first conjugated to protein A Dynabeads (Thermo Fisher). 25 µl of beads were first washed three times with 0.5% BSA in 1x PBS. The beads were then resuspended in 1 ml of 0.5% BSA in 1x PBS, 0.5-2 µl of antibody was added (see Table 2 for antibodies) and incubated for at least 4 h at 4°C with end-over-end rotation to block the beads and bind the antibody. For each IP reaction, 100 µl of concentrated chromatin (corresponding to 1 × 10^6^ cells) was diluted with 900 µl of ChIP dilution buffer (20 mM Tris-Cl pH 8, 150 mM NaCl, 1 mM EDTA, 1% Triton X-100, 0.1 mM PMSF). After washing the antibody:bead conjugates three times with 0.5% BSA in 1x PBS, the diluted chromatin was added and incubated overnight at 4°C with constant rotation. The following morning, the beads were washed once each with low salt wash buffer (20 mM Tris-Cl pH 8, 150 mM NaCl, 2 mM EDTA, 0.1% SDS, 1% Triton X-100), high salt wash buffer (20 mM Tris-Cl pH 8, 500 mM NaCl, 2 mM EDTA, 0.1% SDS, 1% Triton X-100), LiCl buffer (10 mM Tris-Cl pH 8, 250 mM LiCl, 1 mM EDTA, 1% NP40, 1% sodium deoxycholate), and twice with TE buffer (10 mM Tris pH 8, 1 mM EDTA). The washed beads were then resuspended in 100 µl ChIP elution buffer (0.1 M NaHCO_3_, 1% SDS) and shaken vigorously at 1000 rpm in a ThermoMixer for 30 min at 25°C. The beads were centrifuged at 15,000 x *g* for 1 min, magnetized, and the supernatant was taken as the ChIP elution. Input corresponding to 2% of total ChIP reaction and ChIP samples were made up to the same volume and decrosslinked by addition of 200 mM NaCl, 0.3 mg/ml RNase and 0.2 mg/ml proteinase K, followed by incubation for 4 h at 65°C under constant shaking. Decrosslinked DNA was recovered using the Monarch PCR DNA Cleanup Kit (NEB), eluted in 50 µl of elution buffer and analysed by qPCR using a LightCycler 480 instrument (Roche), qPCRBIO SyGreen Blue Mix (PCRBiosystems), and the primers listed in Table 1.

**Table 2.** Antibodies used in this study.

| Antigen | Source | Application |
| --- | --- | --- |
| 6xHis | Sigma H1209 | WB |
| HA | CST 3724 | WB, ChIP (2 µl per ChIP) |
| Tubulin | DHSBAA4-3-5 | WB |
| H3K27ac | CST 8173 | WB, ChIP (0.5 µl per ChIP) |
| H3K9ac | Abcam ab4441 | WB, ChIP (2 µl per ChIP) |
| H2BK5ac | Abcam ab40886 | WB |
| H4K5ac | CST 8647 | WB |
| H2A | CST 3636 | WB |
| H3 | Abcam ab1791 | ChIP (2 µl per ChIP) |

### Reverse transcriptase-qPCR (RT-qPCR)

To extract RNA from ES cells, pellets were resuspended in 1 ml of TriPure reagent (Roche) and purified according to the manufacturer’s protocol. Final RNA pellets were air dried and resuspended in 100 µl RNase-free water. cDNA was prepared from 1 µg of RNA using SuperScript IV reverse transcriptase (Invitrogen). The resulting cDNA was analysed by qPCR as described for ChIP and with the primers listed in Table 1.

## References

An, W., and Roeder, R.G. (2003). Direct association of p300 with unmodified H3 and H4 N termini modulates p300-dependent acetylation and transcription of nucleosomal templates. Journal of Biological Chemistry 278, 1504–1510.

Banka, S., Sayer, R., Breen, C., Barton, S., Pavaine, J., Sheppard, S.E., Bedoukian, E., Skraban, C., Cuddapah, V.A., and Clayton-Smith, J. (2019). Genotype-phenotype specificity in Menke-Hennekam syndrome caused by missense variants in exon 30 or 31 of CREBBP. Am J Med Genet A 179, 1058–1062.

Bannister, A.J., and Kouzarides, T. (1996). The CBP co-activator is a histone acetyltransferase. Nature 384, 641–643.

Barnes, C.E., English, D.M., and Cowley, S.M. (2019). Acetylation & Co: an expanding repertoire of histone acylations regulates chromatin and transcription. Essays Biochem 63, 97–107.

Bhaumik, P., Davis, J., Tropea, J.E., Cherry, S., Johnson, P.F., and Miller, M. (2014). Structural insights into interactions of C/EBP transcriptional activators with the Taz2 domain of p300. Acta crystallographica Section D, Biological crystallography 70, 1914–1921.

Birney, E., Stamatoyannopoulos, J.A., Dutta, A., Guigó, R., Gingeras, T.R., Margulies, E.H., Weng, Z., Snyder, M., Dermitzakis, E.T., Thurman, R.E., et al. (2007). Identification and analysis of functional elements in 1% of the human genome by the ENCODE pilot project. Nature 447, 799–816.

Black, J.B., Adler, A.F., Wang, H.G., D’Ippolito, A.M., Hutchinson, H.A., Reddy, T.E., Pitt, G.S., Leong, K.W., and Gersbach, C.A. (2016). Targeted Epigenetic Remodeling of Endogenous Loci by CRISPR/Cas9-Based Transcriptional Activators Directly Converts Fibroblasts to Neuronal Cells. Cell Stem Cell 19, 406–414.

Bose, D.A., Donahue, G., Reinberg, D., Shiekhattar, R., Bonasio, R., and Berger, S.L. (2017). RNA Binding to CBP Stimulates Histone Acetylation and Transcription. Cell 168, 135-149.e122.

Cheng, A.W., Wang, H., Yang, H., Shi, L., Katz, Y., Theunissen, T.W., Rangarajan, S., Shivalila, C.S., Dadon, D.B., and Jaenisch, R. (2013). Multiplexed activation of endogenous genes by CRISPR-on, an RNA-guided transcriptional activator system. Cell Research 23, 1163–1171.

Conrad, T., Cavalli, F.M.G., Holz, H., Hallacli, E., Kind, J., Ilik, I., Vaquerizas, J.M., Luscombe, N.M., and Akhtar, A. (2012). The MOF chromobarrel domain controls genome-wide H4K16 acetylation and spreading of the MSL complex. Developmental cell 22, 610–624.

Creyghton, M.P., Cheng, A.W., Welstead, G.G., Kooistra, T., Carey, B.W., Steine, E.J., Hanna, J., Lodato, M.a., Frampton, G.M., Sharp, P.a., et al. (2010). Histone H3K27ac separates active from poised enhancers and predicts developmental state. Proceedings of the National Academy of Sciences of the United States of America 107, 21931–21936.

Davey, C.A., Sargent, D.F., Luger, K., Maeder, A.W., and Richmond, T.J. (2002). Solvent mediated interactions in the structure of the nucleosome core particle at 1.9 a resolution. Journal of molecular biology 319, 1097–1113.

Delvecchio, M., Gaucher, J., Aguilar-Gurrieri, C., Ortega, E., and Panne, D. (2013). Structure of the p300 catalytic core and implications for chromatin targeting and HAT regulation. Nature Structural and Molecular Biology 20, 1040–1046.

Filippakopoulos, P., and Knapp, S. (2012). The bromodomain interaction module. FEBS Letters 586, 2692–2704.

Finogenova, K., Bonnet, J., Poepsel, S., Schäfer, I.B., Finkl, K., Schmid, K., Litz, C., Strauss, M., Benda, C., and Müller, J. (2020). Structural basis for decoding active histone methylation marks by Polycomb Repressive Complex 2. bioRxiv.

Heintzman, N.D., Stuart, R.K., Hon, G., Fu, Y., Ching, C.W., Hawkins, R.D., Barrera, L.O., Van Calcar, S., Qu, C., Ching, K.A., et al. (2007). Distinct and predictive chromatin signatures of transcriptional promoters and enhancers in the human genome. Nature genetics 39, 311–318.

Hennekam, R.C.M. (2006). Rubinstein-Taybi syndrome. European Journal of Human Genetics 14, 981–985.

Hilton, I.B., D’Ippolito, A.M., Vockley, C.M., Thakore, P.I., Crawford, G.E., Reddy, T.E., and Gersbach, C.A. (2015). Epigenome editing by a CRISPR-Cas9-based acetyltransferase activates genes from promoters and enhancers. Nature Biotechnology 33, 510–517.

Holbert, M.A., Sikorski, T., Carten, J., Snowflack, D., Hodawadekar, S., and Marmorstein, R. (2007). The human monocytic leukemia zinc finger histone acetyltransferase domain contains DNA-binding activity implicated in chromatin targeting. Journal of Biological Chemistry 282, 36603–36613.

Jefimov, K., Alcaraz, N., Kloet, S.L., Värv, S., Sakya, S.A., Vaagenso, C.D., Vermeulen, M., Aasland, R., and Andersson, a.R. (2018). The GBAF chromatin remodeling complex binds H3K27ac and mediates enhancer transcription. bioRxiv.

Jenkins, L.M.M., Yamaguchi, H., Hayashi, R., Cherry, S., Tropea, J.E., Miller, M., Wlodawer, A., Appella, E., and Mazur, S.J. (2009). Two distinct motifs within the p53 transactivation domain bind to the Taz2 domain of p300 and are differentially affected by phosphorylation. Biochemistry 48, 1244–1255.

Jin, Q., Yu, L.R., Wang, L., Zhang, Z., Kasper, L.H., Lee, J.E., Wang, C., Brindle, P.K., Dent, S.Y.R., and Ge, K. (2011). Distinct roles of GCN5/PCAF-mediated H3K9ac and CBP/p300-mediated H3K18/27ac in nuclear receptor transactivation. EMBO Journal 30, 249–262.

Kabadi, A.M., Ousterout, D.G., Hilton, I.B., and Gersbach, C.A. (2014). Multiplex CRISPR/Cas9-based genome engineering from a single lentiviral vector. Nucleic acids research 42, e147–e147.

Karmodiya, K., Krebs, A.R., Oulad-Abdelghani, M., Kimura, H., and Tora, L. (2012). H3K9 and H3K14 acetylation co-occur at many gene regulatory elements, while H3K14ac marks a subset of inactive inducible promoters in mouse embryonic stem cells. BMC Genomics 13.

Klein, B.J., Jang, S.M., Lachance, C., Mi, W., Lyu, J., Sakuraba, S., Krajewski, K., Wang, W.W., Sidoli, S., Liu, J., et al. (2019). Histone H3K23-specific acetylation by MORF is coupled to H3K14 acylation. Nature Communications 10.

Klein, B.J., Muthurajan, U.M., Lalonde, M.E., Gibson, M.D., Andrews, F.H., Hepler, M., Machida, S., Yan, K., Kurumizaka, H., Poirier, M.G., et al. (2016). Bivalent interaction of the PZP domain of BRPF1 with the nucleosome impacts chromatin dynamics and acetylation. Nucleic Acids Research 44, 472–484.

Kraus, W.L., Manning, E.T., and Kadonaga, J.T. (1999). Biochemical Analysis of Distinct Activation Functions in p300 That Enhance Transcription Initiation with Chromatin Templates. Molecular and Cellular Biology 19, 8123–8135.

Lalonde, M.E., Avvakumov, N., Glass, K.C., Joncas, F.H., Saksouk, N., Holliday, M., Paquet, E., Yan, K., Tong, Q., Klein, B.J., et al. (2013). Exchange of associated factors directs a switch in HBO1 acetyltransferase histone tail specificity. Genes and Development 27, 2009–2024.

Lowary, P.T., and Widom, J. (1998). New DNA sequence rules for high affinity binding to histone octamer and sequence-directed nucleosome positioning. Journal of Molecular Biology 276, 19–42.

Luger, K., Mäder, A.W., Richmond, R.K., Sargent, D.F., and Richmond, T.J. (1997). Crystal structure of the nucleosome core particle at 2.8 A resolution. Nature 389, 251–260.

Marmorstein, R., and Zhou, M.M. (2014). Writers and readers of histone acetylation: Structure, mechanism, and inhibition. Cold Spring Harbor Perspectives in Biology 6.

Menke, L.A., Gardeitchik, T., Hammond, P., Heimdal, K.R., Houge, G., Hufnagel, S.B., Ji, J., Johansson, S., Kant, S.G., Kinning, E., et al. (2018). Further delineation of an entity caused by CREBBP and EP300 mutations but not resembling Rubinstein–Taybi syndrome. American Journal of Medical Genetics, Part A 176, 862–876.

Menke, L.A., van Belzen, M.J., Alders, M., Cristofoli, F., Ehmke, N., Fergelot, P., Foster, A., Gerkes, E.H., Hoffer, M.J.V., Horn, D., et al. (2016). CREBBP mutations in individuals without Rubinstein–Taybi syndrome phenotype. American Journal of Medical Genetics, Part A 170, 2681–2693.

Miller, M., Dauter, Z., Cherry, S., Tropea, J.E., and Wlodawer, A. (2009). Structure of the Taz2 domain of p300: Insights into ligand binding. Acta Crystallographica Section D: Biological Crystallography 65, 1301–1308.

Ogryzko, V.V., Schiltz, R.L., Russanova, V., Howard, B.H., and Nakatani, Y. (1996). The transcriptional coactivators p300 and CBP are histone acetyltransferases. Cell 87, 953–959.

Oike, Y., Takakura, N., Hata, A., Kaname, T., Akizuki, M., Yamaguchi, Y., Yasue, H., Araki, K., Yamamura, K.I., and Suda, T. (1999). Mice homozygous for a truncated form of CREB-binding protein exhibit defects in hematopoiesis and vasculo-angiogenesis. Blood 93, 2771–2779.

Park, S., Stanfield, R.L., Martinez-Yamout, M.A., Dyson, H.J., Wilson, I.A., and Wright, P.E. (2017). Role of the CBP catalytic core in intramolecular SUMOylation and control of histone H3 acetylation. Proceedings of the National Academy of Sciences of the United States of America 114, E5335–E5342.

Pasini, D., Malatesta, M., Jung, H.R., Walfridsson, J., Willer, A., Olsson, L., Skotte, J., Wutz, A., Porse, B., Jensen, O.N., et al. (2010). Characterization of an antagonistic switch between histone H3 lysine 27 methylation and acetylation in the transcriptional regulation of Polycomb group target genes. Nucleic Acids Research 38, 4958–4969.

Petrij, F., Giles, R.H., Dauwerse, H.G., Saris, J.J., Hennekam, R.C., Masuno, M., Tommerup, N., van Ommen, G.J., Goodman, R.H., and Peters, D.J. (1995). Rubinstein-Taybi syndrome caused by mutations in the transcriptional co-activator CBP. Nature 376, 348–351.

Rada-Iglesias, A., Bajpai, R., Swigut, T., Brugmann, S.A., Flynn, R.A., and Wysocka, J. (2011). A unique chromatin signature uncovers early developmental enhancers in humans. Nature 470, 279–283.

Ruthenburg, A.J., Li, H., Patel, D.J., and Allis, C.D. (2007). Multivalent engagement of chromatin modifications by linked binding modules. Nat Rev Mol Cell Biol 8, 983–994.

Sebé-Pedrós, A., De Mendoza, A., Lang, B.F., Degnan, B.M., and Ruiz-Trillo, I. (2011). Unexpected repertoire of metazoan transcription factors in the unicellular holozoan capsaspora owczarzaki. Molecular Biology and Evolution 28, 1241–1254.

Shogren-Knaak, M., Ishii, H., Sun, J.-M., Pazin, M.J., Davie, J.R., and Peterson, C.L. (2006). Histone H4-K16 acetylation controls chromatin structure and protein interactions. Science (New York, NY) 311, 844–847.

Tanaka, Y., Naruse, I., Maekawa, T., Masuya, H., Shiroishi, T., and Ishii, S. (1997). Abnormal skeletal patterning in embryos lacking a single Cbp allele: A partial similarity with Rubinstein-Taybi syndrome. Proceedings of the National Academy of Sciences of the United States of America 94, 10215–10220.

Tie, F., Banerjee, R., Stratton, C.A., Prasad-Sinha, J., Stepanik, V., Zlobin, A., Diaz, M.O., Scacheri, P.C., and Harte, P.J. (2009). CBP-mediated acetylation of histone H3 lysine 27 antagonizes Drosophila Polycomb silencing. Development 136, 3131–3141.

Vermeulen, M., Mulder, K.W., Denissov, S., Pijnappel, W.W.M.P., van Schaik, F.M.a., Varier, R.a., Baltissen, M.P.a., Stunnenberg, H.G., Mann, M., and Timmers, H.T.M. (2007). Selective anchoring of TFIID to nucleosomes by trimethylation of histone H3 lysine 4. Cell 131, 58–69.

Voigt, P., LeRoy, G., Drury, W.J., Zee, B.M., Son, J., Beck, D.B., Young, N.L., Garcia, B.A., and Reinberg, D. (2012). Asymmetrically modified nucleosomes. Cell 151, 181–193.

Wang, Z., Zang, C., Cui, K., Schones, D.E., Barski, A., Peng, W., and Zhao, K. (2009). Genome-wide Mapping of HATs and HDACs Reveals Distinct Functions in Active and Inactive Genes. Cell 138, 1019–1031.

Wang, Z., Zang, C., Rosenfeld, J.A., Schones, D.E., Barski, A., Cuddapah, S., Cui, K., Roh, T.Y., Peng, W., Zhang, M.Q., et al. (2008). Combinatorial patterns of histone acetylations and methylations in the human genome. Nature Genetics 40, 897–903.

Weinert, B.T., Narita, T., Satpathy, S., Srinivasan, B., Hansen, B.K., Schölz, C., Hamilton, W.B., Zucconi, B.E., Wang, W.W., Liu, W.R., et al. (2018). Time-Resolved Analysis Reveals Rapid Dynamics and Broad Scope of the CBP/p300 Acetylome. Cell 174, 231-244.e212.

Wojciak, J.M., Martinez-Yamout, M.A., Dyson, H.J., and Wright, P.E. (2009). Structural basis for recruitment of CBP/p300 coactivators by STAT1 and STAT2 transactivation domains. EMBO Journal 28, 948–958.

Yao, T.P., Oh, S.P., Fuchs, M., Zhou, N.D., Ch’ng, L.E., Newsome, D., Bronson, R.T., Li, E., Livingston, D.M., and Eckner, R. (1998). Gene dosage-dependent embryonic development and proliferation defects in mice lacking the transcriptional integrator p300. Cell 93, 361–372.

Zentner, G.E., and Henikoff, S. (2013). Regulation of nucleosome dynamics by histone modifications. Nature Structural and Molecular Biology 20, 259–266.

Zhang, T., Zhang, Z., Dong, Q., Xiong, J., and Zhu, B. (2020). Histone H3K27 acetylation is dispensable for enhancer activity in mouse embryonic stem cells. Genome Biol 21, 45.

Zhang, Y., Xue, Y., Shi, J., Ahn, J.W., Mi, W., Ali, M., Wang, X., Klein, B.J., Wen, H., Li, W., et al. (2018). The ZZ domain of p300 mediates specificity of the adjacent HAT domain for histone H3. Nature Structural and Molecular Biology 25, 841–849.

